# Neuronal cholesterol synthesis is essential for repair of chronically demyelinated lesions in mice

**DOI:** 10.1101/2021.08.13.456070

**Authors:** Stefan A. Berghoff, Lena Spieth, Ting Sun, Leon Hosang, Constanze Depp, Andrew O. Sasmita, Martina H. Vasileva, Patricia Scholz, Yu Zhao, Dilja Krueger-Burg, Sven Wichert, Euan R Brown, Kyriakos Michael, Klaus-Armin Nave, Stefan Bonn, Francesca Odoardi, Moritz Rossner, Till Ischebeck, Julia M. Edgar, Gesine Saher

**Affiliations:** Department of Neurogenetics, Max Planck Institute of Experimental Medicine, Göttingen, Germany; Institute for Medical Systems Biology, Center for Molecular Neurobiology Hamburg, Hamburg, Germany; Institute for Neuroimmunology and Multiple Sclerosis Research, University Medical Center Göttingen, Göttingen, Germany; Department of Plant Biochemistry, Albrecht-von-Haller-Institute for Plant Sciences and Göttingen Center for Molecular Biosciences (GZMB), University of Göttingen, Göttingen, Germany; Department of Molecular Neurobiology, Max Planck Institute of Experimental Medicine, Göttingen, Germany; Department of Psychiatry and Psychotherapy, University Hospital, LMU Munich, Munich, Germany; School of Engineering and Physical Sciences, Institute of Biological Chemistry, Biophysics and Bioengineering, James Naysmith Building, Heriot Watt University, Edinburgh, UK; Service Unit for Metabolomics and Lipidomics, Göttingen Center for Molecular Biosciences (GZMB), University of Göttingen, Göttingen, Germany; Axo-glial Group, Institute of Infection, Immunity and Inflammation, College of Medical Veterinary and Life Sciences, University of Glasgow, Glasgow, UK

## Abstract

Astrocyte-derived cholesterol supports brain cells under physiological conditions. However, in demyelinating lesions, astrocytes downregulate cholesterol synthesis and the cholesterol that is essential for remyelination has to originate from other cellular sources. Here, we show that repair following acute versus chronic demyelination involves distinct processes. In particular, we found that in chronic myelin disease, when recycling of lipids is often defective, de novo neuronal cholesterol synthesis is critical for regeneration. By gene expression profiling, genetic loss of function experiments and comprehensive phenotyping, we provide evidence that neurons increase cholesterol synthesis in chronic myelin disease models and MS patients. In mouse models, neuronal cholesterol facilitated remyelination specifically by triggering OPC proliferation. Our data contribute to the understanding of disease progression and have implications for therapeutic strategies in MS patients.

## Introduction

During normal brain development, cholesterol is produced locally by de novo synthesis involving neurons, oligodendrocytes, microglia, and astrocytes (Berghoff et al., 2021; Camargo et al., 2012; Fünfschilling et al., 2012; Saher et al., 2005). Neuronal cholesterol is essential for neurite outgrowth and synapse formation during neurogenesis (Fünfschilling et al., 2012; Mauch et al., 2001) but the highest rates of cholesterol synthesis in the brain are achieved by oligodendrocytes during postnatal myelination (Dietschy, 2009). The resulting cholesterol-rich myelin enwraps, shields, and insulates axons to enable rapid conduction of neuronal impulses. Myelin also provides support to axons, potentially by mobilizing oligodendroglial lipids (Kassmann et al., 2007; Saab and Nave, 2017). In the adult brain, cholesterol synthesis is attenuated to low steady-state levels (Dietschy and Turley, 2004).

Destruction of lipid-rich myelin in demyelinating diseases such as multiple sclerosis (MS) likely impairs neuronal function by disrupting the fine-tuned axon-myelin unit (Stassart et al., 2018). Remyelination is considered crucial for limiting axon damage and slowing progressive clinical disability. Statin-mediated inhibition of the cholesterol synthesis pathway impairs remyelination (Miron et al., 2009). Previously, we showed that following an acute demyelinating episode, oligodendrocytes import cholesterol for new myelin membrane synthesis from damaged myelin that has been recycled by phagocytic microglia (Berghoff *et al.*, 2021). In contrast, de novo oligodendroglial cholesterol synthesis contributes to remyelination only following chronic demyelination (Berghoff *et al.*, 2021; Voskuhl et al., 2019). Notably, we and others showed that astrocytes reduce expression of cholesterol synthesis genes following demyelination (Berghoff *et al.*, 2021; Itoh et al., 2018). As astrocytes are considered to support neurons by providing cholesterol in ApoE-containing lipoproteins in the healthy brain (Dietschy, 2009), the lack of this support in the diseased brain contributes to the disruption of CNS cholesterol homeostasis. Neuronal activity leads to OPC proliferation during development and likely also after demyelination (Bacmeister et al., 2020; Gibson et al., 2014; Marisca et al., 2020). However, neuronal responses to myelin degeneration with regard to cholesterol metabolism, and the contribution of neuronal cholesterol to remyelination, remains unknown.

Here, using mice with cell type-specific inactivation of cholesterol synthesis and models of myelin disease, we assess neuronal versus glial cholesterol metabolism. We compare white and grey matter CNS regions and isolated brain cells in the healthy adult brain and during remyelination. We show that active myelin disease is associated with downregulated expression of cholesterol metabolism in neurons. Surprisingly, during chronic myelin disease, neurons increase cholesterol synthesis. Similarly, neurons in MS brain upregulate a gene profile related to cholesterol synthesis and metabolism in non-lesion areas. Finally, neuronal cholesterol synthesis contributes to remyelination following experimental demyelination. Our data support the essential role of cholesterol synthesis in neurons for remyelination, a role that is likely relevant for MS disease progression.

## Results

### Loss of *Fdft1* in neurons alters white matter cholesterol metabolism

In the adult brain, neuronal synthesis as well as horizontal cholesterol transfer from glial cells meets neuronal cholesterol demands. To evaluate neuronal versus glial cholesterol metabolism, we acutely isolated neurons, astrocytes and oligodendrocytes from brain tissue that contained cortex or subcortical white matter of adult mice (Figure1A). The abundance of neuronal mRNA transcripts related to cholesterol metabolism was compared with oligodendrocyte and astrocyte profiles obtained previously (Berghoff *et al.*, 2021). As expected, neurons showed low steady-state expression levels of cholesterol synthesis genes (*Hmgcr, Fdft1, Cyp51, Dhcr24*) compared to oligodendrocytes and astrocytes (Figure 1B, Table S1). In contrast, several gene transcripts related to cholesterol import (*Apobr, Scarb1, Lrp1*), storage (*Soat1*) and brain export (*Cyp46a1*) were higher in relative abundance. To assess the relevance of cell type-specific cholesterol synthesis, we genetically inactivated squalene synthase (SQS, *Fdft1* gene), an essential enzyme of the sterol biosynthesis pathway, in adult oligodendrocytes (OLcKO, Plp1-CreERT2), OPCs (OPCcKO, CSPG4::CreERT2), astrocytes (AcKO, GLAST::CreERT2), or neurons (NcKO, CaMKII-Cre) (Figure 1C, S1A-B) (Berghoff *et al.*, 2021; Fünfschilling *et al.*, 2012; Saher *et al.*, 2005). Comparable to oligodendroglial and astrocyte conditional mutants (Berghoff *et al.*, 2021), loss of cholesterol synthesis in neurons did not affect peripheral serum cholesterol level or body weight (Figure S1C). In an open field test, neuronal, astrocyte or OPC mutants appeared similar to controls (Figure 1D, S1D-E), whereas OLcKO animals showed signs of anxiety, which were enhanced in OPC/OL double mutants (Figure S1F). Notably, these behavioral changes occurred in the absence of overt myelin / oligodendrocyte deficits (Figure S1G), which were also not observed in the other conditional mutants (Berghoff *et al.*, 2021; Fünfschilling *et al.*, 2012).

**Figure 1.**
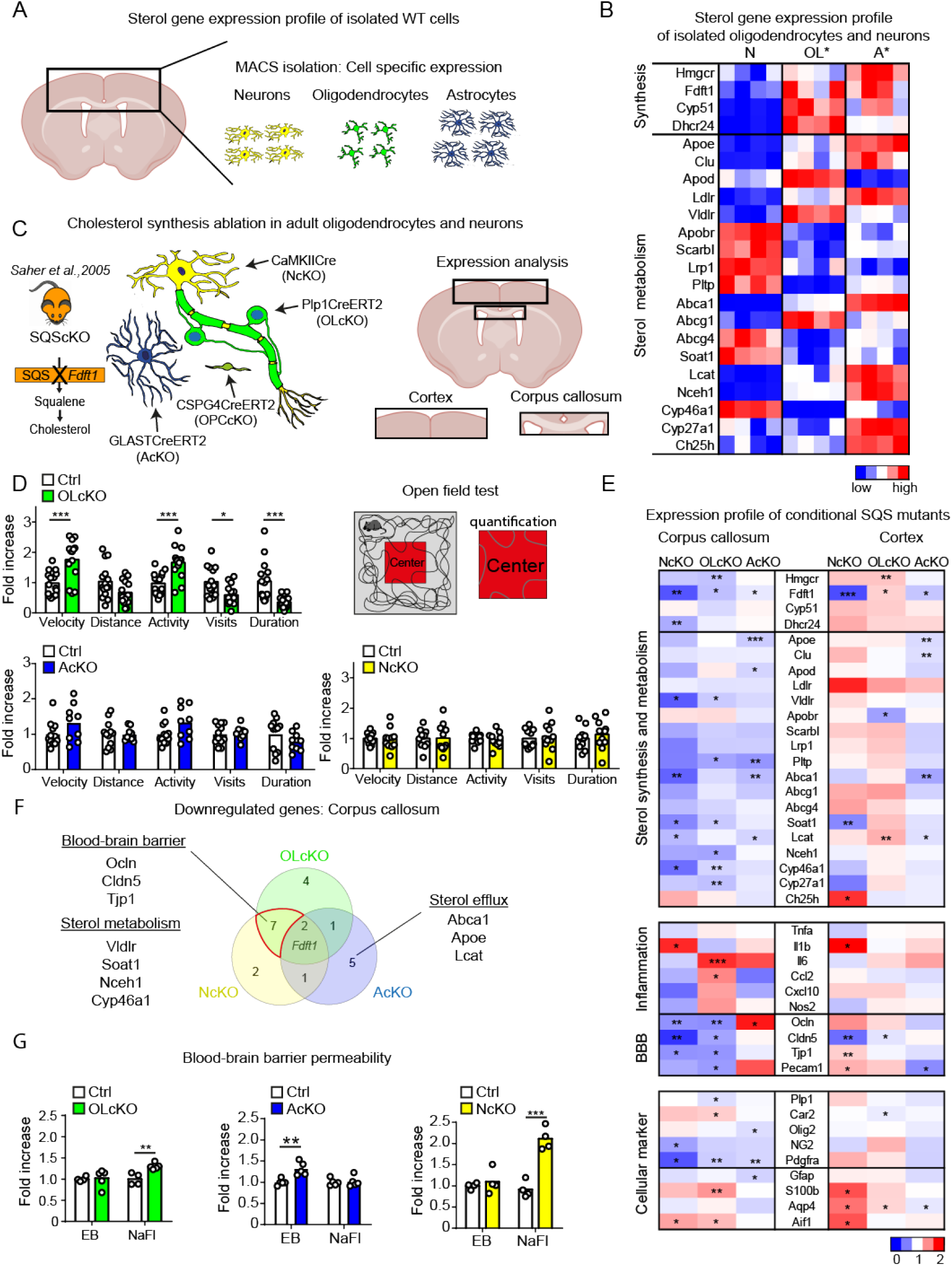
Loss of cholesterol synthesis in oligodendrocytes, astrocytes and neurons. (A) Scheme depicting MACS isolation of neurons, astrocytes and oligodendrocytes from the indicated brain region. (B)Gene expression profile of isolated neurons (N), oligodendrocytes (OL) and astrocytes (A) from adult wild type animals (* Berghoff et al., 2021). (C) Scheme depicting isolation of tissue samples from oligodendrocyte (Plp1-CreERT2), astrocyte (GLAST::CreERT2) and neuronal (CaMKII-Cre) cholesterol synthesis deficient (*Fdft1^fl/fl^*) animals (left) from boxed brain regions for expression analysis. (D) Open field test of OLcKO (n=14), AcKO (n=9) and NcKO (n=8-11) animals compared to corresponding controls (n=10-15) in the center (two-sided Student’s t-test). (E) Expression profile of genes related to cholesterol metabolism, inflammation, blood-brain barrier and cellular identity in indicated CNS tissues of cell type-specific cholesterol synthesis deficient animals. Heat maps show mean fold expression of biological replicates (n=4-7) normalized to controls (two-sided Student’s t-test). (F) Venn diagram of downregulated genes in corpus callosum of animals in (E). (G) Extravasated Evans Blue (EB) and sodium fluorescein (NaFl) in conditional *Fdft1* mutants in oligodendrocytes (OLcKO, n=5), astrocytes (AcKO, n=5) and neurons (NcKO, n=4) compared to corresponding controls (n=4; two-sided Student’s t-test). ***p<0.001, **p<0.01, *p<0.05

Next, we evaluated the impact of conditional loss of squalene synthase / cholesterol synthesis for cholesterol homeostasis by gene transcription profiling of cortex or subcortical white matter (corpus callosum) from the conditional cholesterol synthesis mutants and respective controls. In the cortex, conditional inactivation of cholesterol synthesis in neurons or oligodendrocytes resulted in moderate upregulation of cholesterol synthesis genes, possibly to compensate for the loss of cholesterol synthesis in the affected cell type (Figure 1E, Table S2, S3). In contrast, in all conditional mutants we observed a moderate but consistent downregulation of expression related to cholesterol metabolism in white matter (Figure 1E-F, S1H). Here, reduced expression of genes associated with horizontal cholesterol transfer such as *Abca1*, *Apoe* and *Lcat* was noted in all conditional mutants, particularly in AcKO animals (Figure 1F). In agreement, sterol profiling revealed only moderate alteration in conditional mutants (Figure S2A).

Surprisingly, comparison of significant transcript changes in conditional mutants revealed a marked downregulation of genes related to the blood-brain barrier (BBB) in corpus callosum samples of OLcKO and NcKO mice. This was accompanied by upregulation of few inflammatory mediators with profiles being unique to each mutant (Figure 1E-F). Interestingly, biochemical quantification of BBB permeability revealed that reduced tight junction gene expression was paralleled by increased CNS influx of the small molecular weight BBB tracer NaFl (376 Da) in OLcKO and NcKO brains. Perhaps surprisingly, given the role of astrocytes in BBB formation, this was not the case in AcKO animals (Figure 1F, S1D, SB-C).

In summary, genetic elimination of cholesterol synthesis in oligodendrocytes, neurons and astrocytes leads to altered transcriptional expression of genes related to cholesterol metabolism in grey and white matter. These data confirm that all cell types contribute to cholesterol homeostasis in the adult brain by cell autonomous cholesterol synthesis. Of note, in cortex but also corpus callosum of neuronal *Fdft1* mutants, *Fdft1* expression was significantly reduced and the abundance of cholesterol synthesis intermediates was attenuated (Figure 1E, S2A). This suggests axonal localization of cholesterol synthesis transcripts *in vivo*, contrary to neurons in compartmentalized culture (Vance et al., 2000).

### Myelin disease leads to increased cholesterol synthesis in neurons

Experimental demyelination is associated with axonal pathology (Berghoff et al., 2017b; Nikic et al., 2011), and chronic loss of myelin in multiple sclerosis patients leads to persistent disabilities. Even subtle myelin defects can lead to axonal damage, myelin instability and glial activation, as observed in null mutants of the myelin-specific genes *Plp1* (proteolipid protein 1) and *Cnp1* (2’,3’-cyclic-nucleotide 3’-phosphodiesterase) (Edgar et al., 2009; Edgar et al., 2004; Lappe-Siefke et al., 2003; Trevisiol et al., 2020). To assess neuronal responses to mild alterations in myelin integrity, we combined the well-characterized *Plp1* and *Cnp1* knockout mice with Thy1-EYFPnuc transgenes to label neuronal nuclei, predominantly of callosal projection neurons (CPN) (Wehr et al., 2006). Despite the axonal pathology, CPN loss was not a feature of *Plp1* and *Cnp1* mutants up to 12 months of age as quantified by EYFPnuc+ cell counting (Figure S2D). We then isolated CPN from cortical layer five by fluorescence-directed laser microdissection at various ages (1, 3, 6 and 12 month) for transcriptional profiling (Figure 2A). Neuronal identity of Thy1-EYFPnuc+ cells isolated from cortical layer five was verified by cell type-specific marker gene expression (Figure S2E). Transcriptional profiling revealed 412 differentially expressed genes in neurons from *Cnp1* knockout mice and 104 genes from *Plp1* knockout mice compared to Thy1-EYFPnuc controls (adj. p-val<0.001, Benjamin-Hochberg correction, at least 1.8-fold changed expression) (Figure S2F). Surprisingly, by gene set enrichment analysis (GSEA) the gene set “cholesterol metabolism” was upregulated in CPNs of both mutants (Figure 2B). This included genes involved in cholesterol synthesis (*Hmgcr, Fdft1, Cyp51, Dhcr24*) and transport (*Ldlr*, *Apoe*) (Figure 2C).

**Figure 2.**
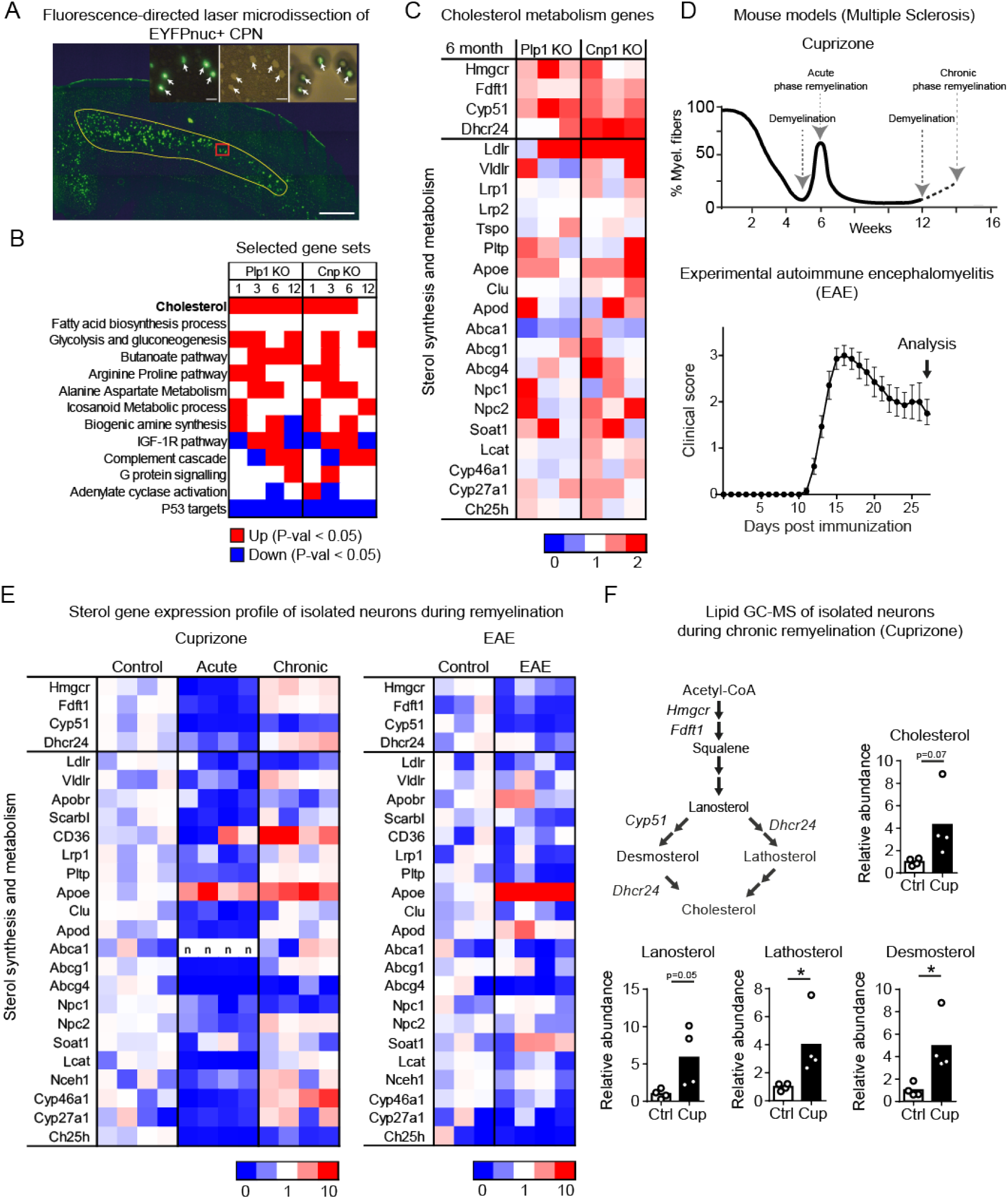
Expression of genes related to cholesterol metabolism in neurons of genetic and experimental myelin disease models. (A) Representative fluorescent micrographs depicting the isolation of EYFPnuc+ callosal projection neurons (CPN) from the primary motor and somatosensory cortex (yellow line) of PLP-KO and CNP-KO mutants (scale 200 μm). Insets show the tissue section before and after laser microdissection, and isolated neurons (scale 20 μm). (B) Selected gene sets in microdissected neurons from PLP-KO and CNP-KO mutants compared to controls (1, 3, 6, 12 months of age). Significantly up-regulated and down-regulated gene sets (P-val<0.05) are indicated in red and blue, respectively. (C) Gene expression profile of genes related to cholesterol metabolism in callosal projection neurons from PLP-KO and CNP-KO mutants at 6 month of age. Heat maps show fold expression of biological replicates (n=3) normalized to controls (Table S5). (D) Time points of analysis related to the course of demyelination/remyelination in the cuprizone model and the clinical score of EAE-induced animals analyzed in (E). (E) Gene expression profile of genes related to cholesterol metabolism in isolated neurons following acute (6 weeks) and chronic (12w+2w) cuprizone challenge (left) and following EAE (right). Heat maps show fold expression normalized to controls (Table S6-S7). Each square represents an individual animal (n=3-4). (F) Cholesterol synthesis pathway with major enzymes and sterol intermediates. Mean relative abundance of sterol intermediates in isolated neurons from chronic cuprizone-treated mice (n=4) compared to untreated controls (n=3), measured by GC-MS (two-sided Student’s t-test).

Considering the possibility that neuronal upregulation of genes related to cholesterol metabolism is a general response to chronic myelin alterations, we analyzed transcriptional profiles of isolated cortical neurons following acute experimental autoimmune encephalomyelitis (EAE) induction and following acute-phase (6 weeks cuprizone) and chronic-phase remyelination (chronic demyelination for 12 weeks followed by two weeks of cuprizone withdrawal, 12+2 weeks) (Figure 2D). During acute disease (EAE or 6 weeks cuprizone), neurons consistently downregulated gene expression related to cholesterol metabolism (Figure 2E). In contrast, during remyelination following chronic demyelination in the cuprizone model, expression of genes involved in cholesterol synthesis (*Hmgcr, Fdft1, Dhcr24*) and transport (*Vldlr*, *Apoe*) were increased (Figure 2E). To test whether the upregulation of cholesterol synthesis genes was functionally relevant, we determined the abundance of cholesterols by GC/MS in the chronic demyelination/remyelination paradigm. Cholesterol as well as several precursors of the cholesterol synthesis pathway were increased 4-5 fold in isolated neurons (Figure 2F), suggesting enhanced cholesterol synthesis in neurons during remyelination in this paradigm.

### Increased cholesterol synthesis gene expression in neurons from MS patients

We next determined whether increased neuronal expression of cholesterol synthesis genes is relevant in human MS and used single-nuclei gene expression profiles from MS patients and healthy control tissue from two recent studies (GSE118257 and GSE124335). We separately merged expression profiles of neurons, oligodendrocytes and astrocytes as annotated in each study (Figure 3A-B, S3A-C). Although the disease history of the MS tissue samples is unknown, we categorized MS samples in two subsets. One comprises the different stages of active MS lesions (termed “lesion”). The other subset (termed “non-lesion”) contains normal-appearing MS tissue that was derived from areas adjacent to active lesions. Neuronal nuclei contributed considerably to each of the MS subsets (3122 lesion, 3547 non-lesion). Next, we performed pairwise comparisons of expression profiles separately, in neurons, oligodendrocytes or astrocytes, focusing on cholesterol metabolism.

**Figure 3.**
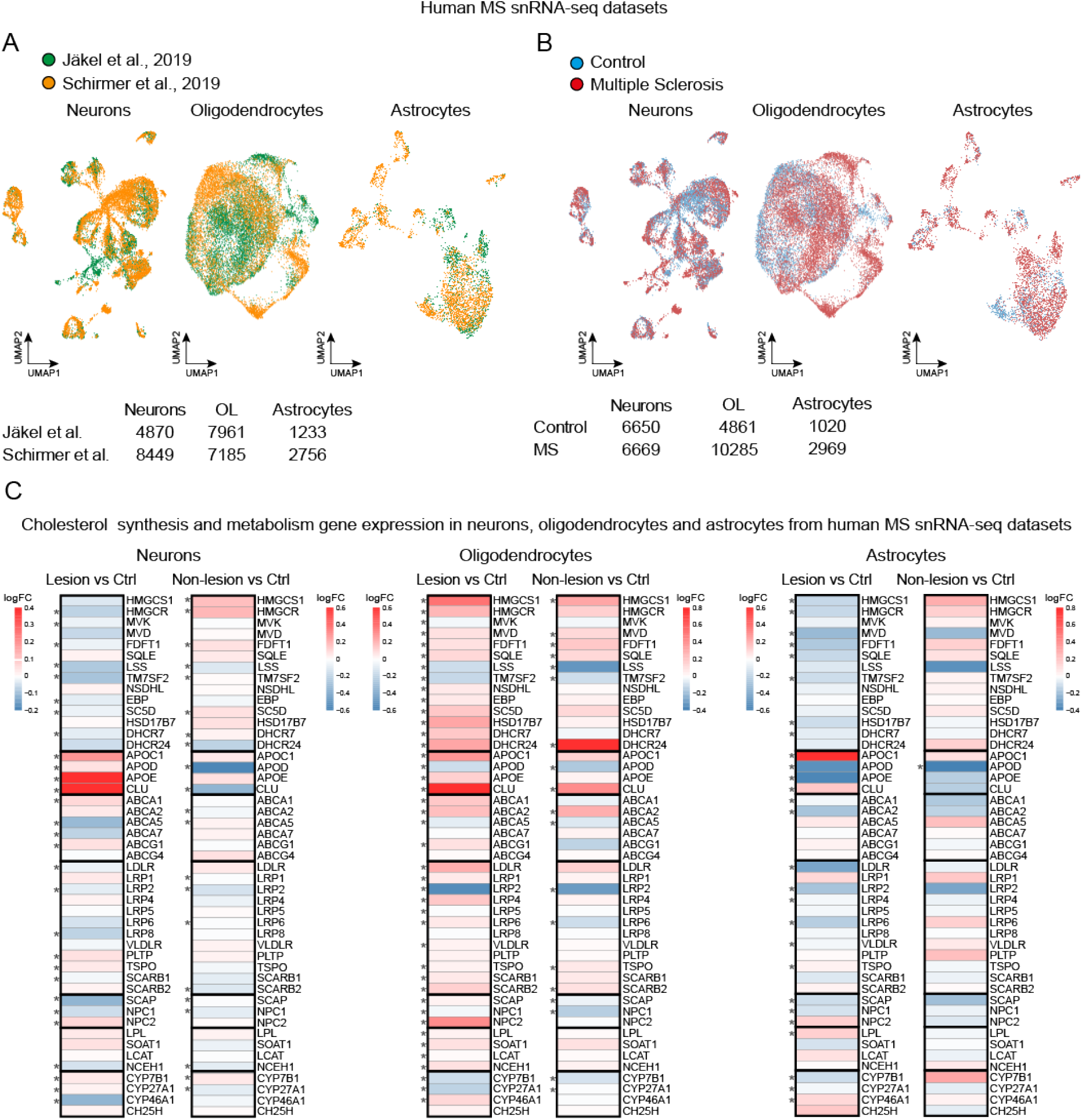
Gene expression related to cholesterol metabolism in neurons, oligodendrocytes and astrocytes from patients with MS. (A) UMAP (Uniform Manifold Approximation and Projection) of neurons, oligodendrocytes and astrocytes from human MS snRNAseq datasets according to dataset (GSE118257, GSE124335). (B) UMAP projection of neurons, oligodendrocytes and astrocytes from human MS snRNAseq datasets according to patient samples identity (GSE118257, GSE124335). (C) Heat map of mean gene expression related to cholesterol metabolism in cellular subsets comparing control and MS (lesion and non-lesion) samples. Data represent log2fold changes (logFC) and p-value (Wilcoxon Rank Sum test, two sided).

As expected, lesion-derived astrocytes showed significantly reduced transcript levels of several genes related to cholesterol synthesis (*HMGCR, FDFT1, DHCR7, DHCR24*), while this gene set was not differentially regulated in non-lesion-derived astrocytes (Figure 3C). In contrast to astrocytes, both oligodendrocytes and neurons in MS lesions, increased expression of apolipoproteins including *APOE*, indicating active participation in local lipid transport in areas of active disease. Moreover, MS oligodendrocytes upregulated genes associated with cholesterol synthesis and metabolism, most markedly in lesions. This confirms the relevance of cholesterol availability for oligodendrocytes in myelin disease (Berghoff *et al.*, 2017b; Berghoff *et al.*, 2021; Voskuhl *et al.*, 2019) and potentially reflects ongoing remyelination or attempts to remyelinate. Notably, neurons in MS lesions showed reduced transcript levels of genes associated with cholesterol synthesis including the rate-limiting enzyme of this process, *HMGCR* (Figure 3C). In contrast, in non-lesion MS tissue, neurons upregulated this gene set, suggesting increased neuronal cholesterol synthesis in normal-appearing MS tissue areas.

### Neuronal *Fdft1* ablation impairs remyelination following chronic cuprizone

The contrasting expression of neuronal cholesterol synthesis genes in lesion versus non-lesion areas of MS brain and in acute versus chronic-phase remyelination in mouse models prompted us to test the importance of this finding for lesion repair. To explore whether loss of neuronal cholesterol synthesis affects remyelination efficiency, NcKO animals were challenged with acute and chronic demyelination paradigms using EAE and cuprizone. We evaluated disease expression, remyelination (Gallyas), oligodendrocyte differentiation (CAII), number of oligodendrocyte linage cells (Olig2) and gliosis (GFAP, Iba1, MAC3).

As anticipated, loss of neuronal cholesterol synthesis did not affect acute-phase remyelination in cuprizone treated mice (Figure 4A-B, S4A-C, Video1) or pathology following immune mediated myelin degeneration (Figure 4C-E). However, after cuprizone-induced chronic demyelination, we observed reduced oligodendrocyte density and impaired remyelination in both the corpus callosum and cortex of NcKO animals compared to controls (Figure 4F-H, S4A-C). Defective repair of chronically demyelinated lesions in NcKO animals occurred without affecting gliosis, neuronal degeneration (NeuN+ cell number, Fluorojade, TUNEL) or axonal stress (APP+ spheroids) (Figure S4D-F). The degree of sustained hypomyelination in NcKO animals was comparable to mutants with inactivated cholesterol synthesis in OPCs (NcKO 53±5% of controls, OPCcKO 55±3%; Figure 4I). Further, both conditional mutants displayed comparable defects in motor performance after chronic cuprizone administration (Figure S4G). However, in contrast to oligodendroglial mutants that showed normal OPC densities, the loss of neuronal cholesterol synthesis caused a reduction of Olig2/PCNA+ proliferating OPCs in the corpus callosum (Figure 4G, I, J). Interestingly, cholesterol administration enhanced OPC proliferation and differentiation in myelinating co-cultures, but only OPC proliferation was impaired by abolishing neuronal activity (Figure 4K, S4H). Together, these findings raise the possibility that elevating neuronal cholesterol synthesis is essential for OPC proliferation and differentiation to facilitate remyelination.

**Figure 4.**
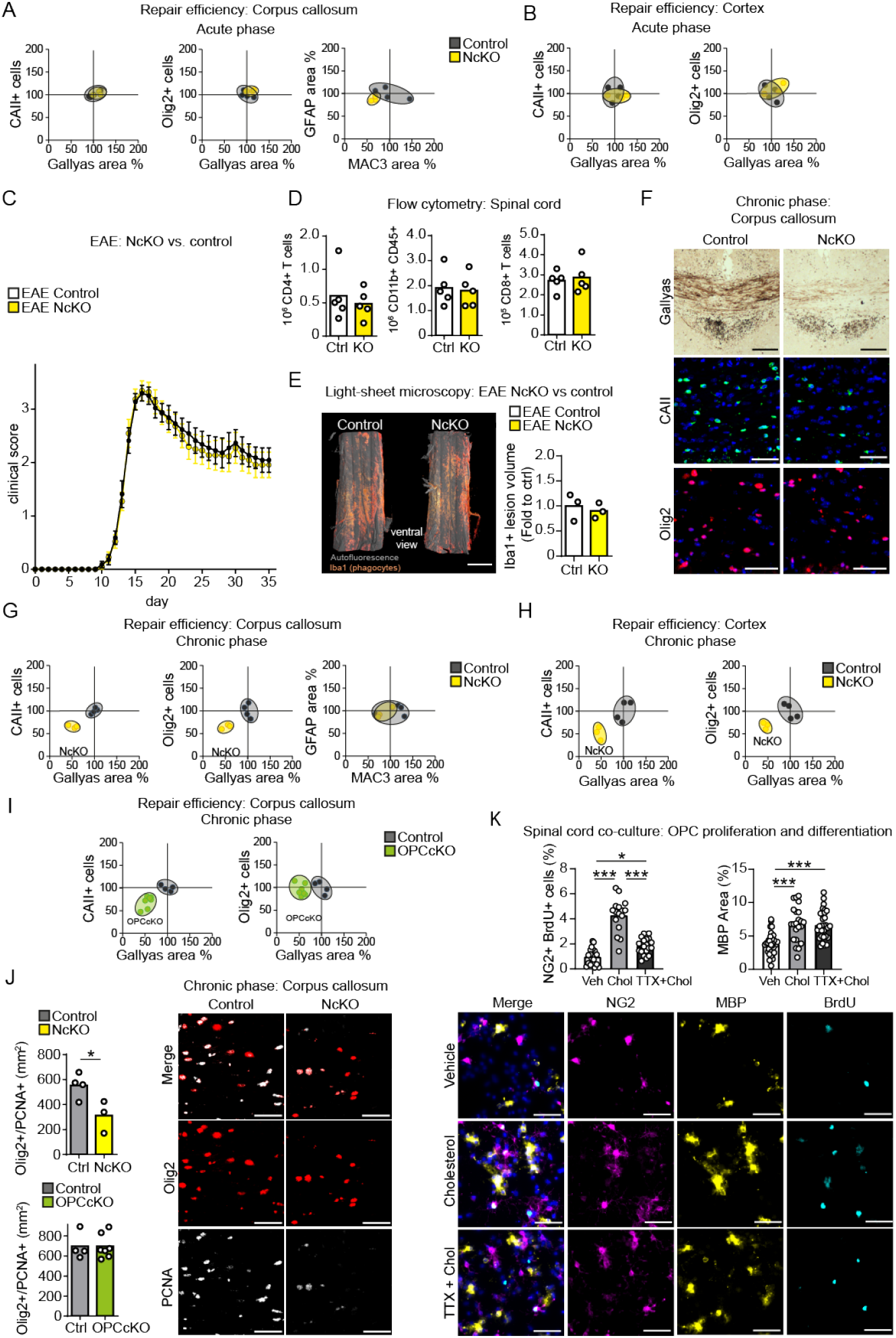
Neuronal *Fdft1* ablation impairs remyelination following chronic cuprizone. (A-B) Repair efficiency and gliosis in the corpus callosum (A) and cortex (B) during acute cuprizone (6 weeks) in NcKO (n=4 animals) compared to controls (n=4, set to 100%) based on histochemical stainings for myelin (Gallyas), oligodendrocytes (CAII), oligodendrocyte lineage cells (Olig2), microgliosis (MAC3) and astrogliosis (GFAP), shown as 95% confidence ellipses with individual data points. (C) Mean clinical EAE score ± SEM of control and NcKO mice (n=11). (D) Flow cytometric quantification of inflammatory cells of EAE animals in (C, n=5). (E) Representative light sheet microcopy with quantification of lumbar spinal cord from NcKO and controls 35d after EAE induction, stained for Iba1+ phagocytes (n=3). (F) Representative micrographs of the corpus callosum of control and NcKO animals following chronic cuprizone challenge (12w+2w) illustrating myelination (Gallyas), oligodendrocytes (CAII), and oligodendrocyte lineage cells (Olig2). (G-H) Repair efficiency and gliosis in the corpus callosum (G) and cortex (H) during chronic cuprizone (12w+2w) in NcKO mice compared to controls (n=4, set to 100%). (I) Repair efficiency in the corpus callosum during chronic cuprizone (12w+2w) in OPC cholesterol mutants (OPCcKO, n=7) compared to controls (n=4, set to 100%). (J) Representative micrographs of Olig2/PCNA double labeling in the corpus callosum with quantification of NcKO (n=3) and OPCcKO (n=7) mutants compared to controls (n=4) following chronic remyelination (12w+2w; two-sided Student’s t-test). **p<0.01, *p<0.05, scales 50 μm, 1 mm (E). (K) Mean OPC proliferation (NG2+, BrdU+) and oligodendrocyte differentiation (MBP+) in spinal cord cultures at 17 days *in vitro* (n=16-31 images from 2-3 cultures) in the presence of cholesterol (10 μg/ml) with or without inhibition of neuronal activity by TTX (1μM). Data show the mean number of NG2+/BrdU+ positive cells normalized to DAPI+ cell density and the mean MBP-positive area with individual values (one-way ANOVA with Sidak’s post-test; Scale 50 μm).

## Discussion

Complete functional recovery from demyelinating episodes and prevention of persistent disabilities is the ultimate goal of multiple sclerosis (MS) therapies. In addition to decreasing the rate of demyelinating events and dampening pathological inflammation, the support of remyelination has come into focus in MS drug development (Plemel et al., 2017). Repair following immune-mediated degeneration of oligodendrocytes and destruction of myelin involves the resolution of inflammation, OPC migration, proliferation and differentiation (Reich et al., 2018). Tissue regeneration is achieved by synthesis of lipid- and cholesterol-rich myelin membranes by newly differentiated oligodendrocytes and oligodendrocytes that survived the immune attack (Franklin et al., 2020). Remyelination contributes to neuroprotection, the restoration of impulse conduction and may facilitate (re)establishing neural circuits (Bacmeister *et al.*, 2020).

### Disparate repair processes during acute and chronic myelin disease

With respect to lipid metabolism, repair after an acute demyelinating episode differs markedly from repair in chronic myelin disease or after repeated demyelinating events. Tissue remodeling and repair after an acute demyelinating attack is coordinated by microglial/macrophage activation and lipid recycling (Berghoff *et al.*, 2021; Cunha et al., 2020; Miron et al., 2013). Correspondingly, acute-phase remyelination is independent of squalene synthase / *Fdft1* inactivation in oligodendrocytes, OPCs, astrocytes, or neurons (this study and Berghoff *et al.*, 2021).

In chronic myelin disease, myelin debris is largely cleared from demyelinated lesions. Correspondingly, in chronically demyelinated lesions in cuprizone-treated mice (Berghoff *et al.*, 2017b) and in chronically inactive lesions in MS patients (Hess et al., 2020) predominantly lipid-laden foamy microglia/macrophages remain. This suggests that lipid recycling by microglia could be inefficient in chronically demyelinated lesions. These lipid trafficking defects in microglia could not only impede remyelination but also aggravate axonal damage, as observed in TREM2 mutants (Cantoni et al., 2015).

As cholesterol is essential for myelin formation (Saher *et al.*, 2005), these findings underscore the necessity of local de novo synthesis. This is supported by the observation that statin-mediated inhibition of cholesterol synthesis blocks remyelination (Miron *et al.*, 2009). However, astrocytes, which provide cholesterol in the healthy brain, fail as a local cholesterol source in myelin disease as they strongly downregulate its synthesis both in mouse models (Berghoff *et al.*, 2021; Borggrewe et al., 2021; Itoh *et al.*, 2018) and MS lesions (this study). We hypothesize that chronic myelin disease depletes neuronal and oligodendroglial cholesterol levels, which triggers cell-autonomous cholesterol synthesis by feedback regulation. In agreement, the cholesterol for remyelination originates at least partially from de novo oligodendroglial synthesis (Berghoff *et al.*, 2021; Jurevics et al., 2002; Voskuhl *et al.*, 2019). The finding that dietary cholesterol supplementation also supports myelin repair (Berghoff *et al.*, 2017b; Saher et al., 2012) suggests that endogenous cholesterol synthesis is insufficient for complete remyelination. In the current study, we provide evidence that neuronal cholesterol synthesis is essential for repair of chronically demyelinated lesions.

### Neuronal cholesterol synthesis during remyelination

Following chronic demyelination, we found impaired remyelination in both white and grey matter of animals lacking neuronal cholesterol synthesis. This is in agreement with the increased neuronal expression of cholesterol synthesis genes in human non-lesion MS tissue and following chronic experimental demyelination. In neuronal mutants of cholesterol synthesis, but not in corresponding oligodendroglial mutants, we showed a marked reduction in the density of oligodendrocyte lineage cells. This finding points to an inference with repair that precedes myelin membrane synthesis, likely in OPC proliferation and oligodendrocyte differentiation.

What is the mechanism by which regeneration of chronically demyelinated white matter tracts such as the corpus callosum benefits from neuronal cholesterol synthesis? Neuronal electrical activity triggers OPC proliferation and oligodendrocyte differentiation during development and after demyelination through unknown signals (Demerens et al., 1996; Marisca *et al.*, 2020; Mitew et al., 2018; Ortiz et al., 2019). The hyperactivity of cortical neurons observed after acute demyelination (Bacmeister *et al.*, 2020) could offset conduction deficits of demyelinated white matter tracts (Crawford et al., 2009). This could lead to synaptic vesicle release from callosal axons (Almeida et al., 2020; Pfeiffer et al., 2019) and provide the endogenous signal to white matter OPCs to proliferate and initiate repair. We observed that neuronal cholesterol plays a role in this process. In cultured OPCs, short term pharmacological inhibition of cholesterol synthesis induces OPC differentiation (Miron et al., 2007). It is possible that reducing cellular cholesterol levels in OPCs decreases the probability of receiving neuronal input via neurotransmitter channels (Korinek et al., 2020). In agreement, we showed that administration of cholesterol facilitates OPC proliferation (this study). However, OPC proliferation was amplified only, when cholesterol was supplied in the context of neuronal activity. It is possible that neurotransmitter release occurs concomitant with release of neuronal cholesterol. This could be a means to prevent (premature) OPC differentiation and to generate sufficient numbers of oligodendrocyte lineage cells to accomplish repair.

In addition, cholesterol-depleted denuded axons as in chronic lesions of NcKO mice are likely more fragile as suggested from increased plasma membrane tether forces of cholesterol synthesis mutant neurons (Fünfschilling *et al.*, 2012). Especially when electrically hyperactive, impaired stability of axonal membranes could increase axon damage and the probability of conduction blocks. Moreover, cholesterol is essential for the biogenesis and exocytosis of synaptic vesicles (Linetti et al., 2010; Thiele et al., 2000). This is compatible with the observation that cholesterol synthesis deficient neurons show reduced spontaneous activity in culture (Fünfschilling *et al.*, 2012).

Several studies have highlighted the importance of cholesterol availability for developmental OPC proliferation, oligodendrocyte differentiation and myelin membrane synthesis (Mathews and Appel, 2016; Saher *et al.*, 2012; Zhao et al., 2016). The finding that administration of dietary cholesterol during chronic-phase remyelination increases the density of proliferating OPCs (Berghoff *et al.*, 2017b) suggests that the entire oligodendrocyte lineage can benefit from externally administered cholesterol. It is possible that neurons increase cholesterol production not only to fulfill their own cholesterol demands but also to support oligodendrocytes to synthesize myelin. Indeed, cholesterol synthesis in callosal axons would position the lipid optimally to supply proliferating OPCs and newly differentiating oligodendrocytes for remyelination. This suggestion is reinforced by the observation that electrically hyperactive neurons in demyelinated lesions release lipids as ApoE-containing lipoproteins (Ioannou et al., 2019; Xu et al., 2006). Our data show that increased expression of neuronal cholesterol synthesis genes is paralleled by strongly elevated expression of ApoE. Thus, following chronic demyelination, neurons could export cholesterol via ApoE to support myelination by oligodendrocytes in a lipid-poor environment. Although we have not measured neuronal activity in demyelinated conditional mutants, we speculate that *Fdft1* mutant neurons produce less activity-dependent pro-repair signals to local OPCs than controls.

Taken together, our data show that loss of neuronal cholesterol synthesis strongly impairs remyelination with relevance for human MS disease. Our study confirms distinct cell type-specific roles in brain cholesterol biogenesis and import during remyelination and provides an additional explanation for disease progression related to age-associated decline of cholesterol synthesis (Berghoff *et al.*, 2021; Scalfari, 2019; Thelen et al., 2006). Further studies are needed to design therapeutic strategies that stimulate cholesterol synthesis in affected neuronal populations.

## Materials and Methods

### Animals

All animal studies were performed in compliance with the animal policies of the Max Planck Institute of Experimental Medicine, and were approved by the German Federal State of Lower Saxony. Animals were group-housed (3-5 mice) with 12 hour dark/light cycle and had access to food and water ad libitum. Adult male and female C57BL/6N mice (8-10 weeks of age) or cholesterol synthesis mutants (8–10 weeks of age) were taken for all experiments. Male mice were subjected to cuprizone experiments. Female mice were used for non-induced pathology experiments. Animals of same gender were randomly assigned to experimental groups (3-12 mice). Cholesterol synthesis mutants in this study were generated by crossbreeding cell type-specific Cre-driver lines (see Key Resources table) with mice harboring squalene synthase floxed mice (*Fdft1^flox/flox^*). Conditional mutants were compared with the respective Cre or homozygous floxed controls, i.e. CaMKII-Cre::*Fdft1^flox/flox^* mutants and *Fdft1^flox/flox^* controls, Plp1-CreERT2::*Fdft1^flox/flox^* mutants and *Fdft1^flox/flox^* controls, Cspg4/NG2^CreERT2/+^::*Fdft1^flox/flox^* and Cspg4/NG2^CreERT2/+^ controls, GLASTCreERT2/+::*Fdft1flox/flox* mutants and GLASTCreERT2/+ controls, Cspg4/NG2^CreERT2/+^::Plp1-CreERT2::*Fdft1^flox/flox^* mutants and Plp1-CreERT2::*Fdft1^flox/flox^* and *Fdft1*^*flox/flox*^ controls. *Cnp* null and *Plp1* null mice were crossbreed to Thy1-EYFPnuc mice to generate CNP mutants (TYNC +/−, *Cnp* −/−, *Plp1* +/y) and PLP mutants (TYNC +/−, *Cnp* +/+, *Plp1* −/y) that were compared with TYNC +/− mice.

### Tamoxifen induced recombination

Transgenic mice received tamoxifen either by oral administration, three times every second day at a concentration of 0.4 mg/g body weight dissolved in corn oil:ethanol (1:9) or by intraperitoneal injections on 5 consecutive days at a concentration of 75 μg/g body weight.

### Experimental autoimmune encephalomyelitis (EAE)

MOG-EAE was induced by immunizing subcutaneously with 200 mg myelin oligodendrocyte glycoprotein peptide 35–55 (MOG35–55) in complete Freund’s adjuvant (M. tuberculosis at 3.75 mg ml-1) and i.p. injection twice with 500 ng pertussis toxin as described (Berghoff *et al.*, 2017b; Berghoff *et al.*, 2021). Animals were examined daily and scored for clinical signs of the disease. If disease did not start within 15 days after induction or the clinical score rose above 4, animals were excluded from the analysis. The clinical score was: 0 normal; 0.5 loss of tail tip tone; 1 loss of tail tone; 1.5 ataxia, mild walking deficits (slip off the grid); 2 mild hind limb weakness, severe gait ataxia, twist of the tail causes rotation of the whole body; 2.5 moderate hind limb weakness, cannot grip the grid with hind paw, but able to stay on a upright tilted grid; 3 mild paraparesis, falls down from a upright tiled grid; 3.5 paraparesis of hind limbs (legs strongly affected, but move clearly); 4 paralysis of hind limbs, weakness in forelimbs; 4.5 forelimbs paralyzed; 5 moribund/dead.

### Cuprizone

Cuprizone pathology was induced by feeding mice with 0.2% w/w cuprizone (Sigma-Aldrich) in powder chow. Mice received cuprizone for ‘acute remyelination’ (6 weeks) and ‘chronic remyelination’ (12 weeks followed by 2 weeks normal chow) paradigms. Chow was replaced three times a week. Age-matched untreated controls were fed powder chow without cuprizone.

### Serum Analysis

Blood was collected by cardiac puncture, and serum was prepared after 4h clotting by centrifugation. Cholesterol measurements were done with the Architect II system (Abbott Diagnostics).

### Open field

Exploratory activity in a novel environment was tested in an open field chamber (50×50×50 cm) at 20 lux light intensity. Individual female mice at the age of 22 weeks were placed into left bottom corner of the open field chamber. The exploratory behavior of the mouse was recorded for 10 min using an overhead camera system and scored automatically using the Viewer software (Biobserve, St. Augustin, Germany). The overall traveled distance was analyzed as a parameter of general activity. Time, distance and visits in the center area (25×25 cm) was analyzed to measure behavior related to anxiety. Results were normalized to the mean of corresponding control animals and statistically analyzed using one-way ANOVA with Sidak’s post-test.

### Blood-brain barrier permeability

Measurements of BBB permeability were done as described (Berghoff et al., 2017a). Tracers were i.v. injected (Evans Blue 50 mg/g body weight, sodium fluorescein 200 mg/g body weight). Animals were flushed with PBS. Brain samples were isolated, weighed, lyophilized at a shelf temperature of –56 °C for 24h under vacuum of 0.2 mBar (Christ LMC-1 BETA 1-16), and extracted with formamide at 57°C for 24h on a shaker at 300 rpm. Integrated density of tracer fluorescence was determined in triplicates after 1:3 ethanol dilutions to increase sensitivity. Tracer concentration was calculated using a standard curve prepared from tracer spiked brain samples.

### Flow cytometry

Single-cell suspensions from spinal cord were obtained via mechanical dissociation on a cell strainer. Immune cells were separated over a two-phase Percoll-density gradient by centrifugation. Staining of CD4+ T cells, CD8+ T cells and CD45/CD11b+ cells (macrophages/microglia) was performed using the following antibodies in a 1:200 dilution: anti-CD3e (clone 145-2C11, BioLegend), anti-CD4 (clone GK 1.5, BD), anti-CD8 (clone 53-6.7, BD), anti-CD11b (clone M1/70, BioLegend), anti-CD45.2 (clone 104, BioLegend). The addition of CaliBRITE APC beads (BD) allowed for cell quantification. Flow cytometry was performed using a CytoFLEX S (Beckman Coulter) operated by CytExpert software (Beckman Coulter, v2.4).

### Magnetic cell isolation (MACS)

Glial cells and neurons were isolated according to the adult brain dissociation protocol (Miltenyi biotec) form corpus callosum and/or cortex. Antibody labeling was done according to the Microbead kit protocols (Miltenyi biotec) for oligodendrocytes (O4) or astrocytes (ACSA-2). Neurons were isolated by negative selection. Purity of cell populations was routinely determined by RT-qPCR on extracted and reverse transcribed RNA.

### Expression analyses

For expression analyses of tissue samples, mice were killed by cervical dislocation. Samples were quickly cooled and region of interest prepared. RNA was extracted using RNeasy Mini kit (Qiagen). cDNA was synthesized with Superscript III (Invitrogen). Concentration and quality of RNA was evaluated using a NanoDrop spectrophotometer and RNA Nano (Agilent). RNA from MACS-purified cells was extracted using QIAshredder and RNeasy protocols (Qiagen). cDNA was amplified by Single Primer Isothermal Amplification (Ribo-SPIA® technology) using Ovation PicoSL WTA System V2 (NuGEN) following the manufactures protocol. Quantitative PCRs were done in triplicates using the GoTaq qPCR Master Mix (Promega, A6002) and the LightCycler 480 Instrument (Roche Diagnostics). Expression values were normalized to the mean of housekeeping genes. Quantification was done by applying the ΔΔCt method, normalized to experimental controls (set to 1). All primers (Table S1) were designed to fulfill optimal criteria e.g. length (18-22 bp), melting temperature (52-58°C), GC content (40-60%), low number of repeats, and amplicon length (<220 bp). All primers were intron-spanning.

### Histochemistry

Mice were perfused with 4% formaldehyde (PFA). In case of cuprizone treated animals, brain samples were cut at Bregma 1.58 to account for regional specificity of cuprizone pathology. Tissue was postfixed overnight, embedded in paraffin and cut into 5 μm sections (HMP 110, MICROM). For Gallyas silver impregnation, deparaffinized sections were incubated with a 2:1 mixture of pyridine and acetic anhydride for 30 min at room temperature (RT) to minimize background and increase myelin. Tissue was washed with ddH_2_0, following heating in incubation solution (0.1% [w/v] ammonium nitrate, 0.1% [w/v] silver nitrate, 12‰ [w/v] sodium hydroxide pH 7.5) for 1 min (100 W) and further incubation for 10 min at RT. After washing with 0.5% [v/v] acetic acid three times for 5 min, sections were incubated in developer solution for 3-10 min. For reconstitution of the developer, 70 ml of solution B (0.2% [w/v] ammonium nitrate, 0.2% [w/v] silver nitrate, 1% [w/v] wolframosilicic acid) was added to 100 ml of solution A (5% [w/v] sodium carbonate) with constant and gentle shaking and then slowly added to 30ml solution C (0.2% [w/v] ammonium nitrate, 0.2% [w/v] silver nitrate, 1% [w/v] wolframosilicic acid, 0.26% [w/v] PFA). The reaction was stopped and fixed by washing in 1.0% [v/v] acetic acid and 2% [v/v] sodium thiosulfate. Tissue was dehydrated and mounted using Eukitt. For detection of apoptotic cells, a TUNEL assay was done according to the manufacturer (Promega G7130). Fluoro-Jade C staining (Sigma, AG 325) was done according to the manufacturers’ instructions. Immunohistological stainings were done on deparaffinized sections followed by antigen-retrieval in sodium citrate buffer (0.01 M, pH 6.0). For chromogenic stainings, blocking of endogenous peroxidase activity with 3% hydrogen peroxide was performed followed by 20% goat serum block and incubation with primary antibodies. Detection was carried out with the LSAB2 System-HRP (anti-rabbit/mouse LSAB2 Kit Dako Cat#K0679, dilution 1:100) or the VECTASTAIN Elite ABC HRP Kit (Vector Labs, Anti-Rat IgG Vector Cat#BA-9400, dilution 1:100). HRP substrate 3,30-diaminobenzidine (DAB) was applied by using the DAB Zytomed Kit (Zytomed Systems GmbH). Nuclear labeling was done by hematoxylin stain. For immunofluorescence detection, blocking was performed with serum-free protein block (Dako).

Primary antibodies were diluted in 2% bovine serum albumin (BSA)/PBS and incubated for 48 h followed by incubation with fluorophore-coupled secondary antibodies (Alexa488 donkey anti-mouse Invitrogen Cat #A-21202, dilution 1:1000; Alexa488 donkey anti-rabbit Invitrogen Cat #A-21206,dilution 1:1000; Alexa555 donkey anti-rabbit Invitrogen Cat #A-31572, dilution 1:1000). Stained sections were analyzed on an Axio Imager.Z1 (Zeiss) equipped with an AxioCam MRc3, x0.63 Camera Adaptor and the ZEN 2012 blue edition software using 10x objective (Plan Apochromat ×10/0.45 M27) or 20x objective (Plan-Apochromat x20/0.8) and evaluated with Image J software. Quantification of areas (Gallyas, GFAP, MAC3) were done by applying semi-automated ImageJ software macro including thresholding and color deconvolution. Two to four sections per animal were analyzed.

### Quantification of sterols

Sterol abundance was quantified by lipid gas chromatography coupled to mass spectrometry (GC-MS) in acutely isolated neurons and tissue samples (4-5 animals grouped for each replicate). Samples were lyophilized at a shelf temperature of –56 °C for 24 h under vacuum of 0.2 millibars (Christ LMC-1 BETA 1-16) and weighed for the calculation of water content and normalization. Metabolites were extracted in a two-phase system of 3:1 methyl-tert-butyl ether:methanol (vol/vol) and water, and pentadecanoic acid was added as an internal standard. The organic phase (10–200 μl) was dried under a stream of nitrogen, dissolved in 10–15 μl pyridine and derivatized with twice the volume of N-methyl-N-(trimethylsilyl) trifluoroacetamide (MSTFA) to transform the sterols and the standard to their trimethylsilyl (TMS) derivatives. Each sample was analyzed twice, with a higher split to quantify cholesterol and with a lower split to measure all other sterols. The samples were analyzed on an Agilent 5977N mass-selective detector connected to an Agilent 7890B gas chromatograph equipped with a capillary HP5-MS column (30 m × 0.25 mm; 0.25-μm coating thickness; J&W Scientific, Agilent). Helium was used as a carrier gas (1 ml/min). The inlet temperature was set to 280 °C, and the temperature gradient applied was 180°C for 1 min, 180–320°C at 5 K min–1 and 320°C for 5 min. Electron energy of 70 eV, an ion source temperature of 230 °C and a transfer line temperature of 280°C were used. Spectra were recorded in the range of 70–600 Da/e (ChemStation Software D.01.02.16). Sterols were identified by the use of external standards.

### Light sheet microscopy

PFA immersion fixed spinal cord segments were processed for whole mount immune-labelling and tissue clearing following a modified iDISCO protocol (Berghoff *et al.*, 2021). Briefly, samples were dehydrated in ascending methanol (MeOH)/PBS series followed by overnight bleaching /permeabilization in a mix of 5% H_2_O_2_/20% DMSO/MeOH at 4°C. Samples were further washed in MeOH and incubated in 20% DMSO/MeOH at RT for 2h. Then, samples were rehydrated using a descending methanol/PBS series and further washed with in PBS/0.2% TritonX-100 for 2h. The samples were then incubated overnight in 0.2% TritonX-100, 20% DMSO, and 0.3 M glycine in PBS at 37°C and blocked using PBS containing 6% goat serum, 10% DMSO and 0.2% Triton-X100 for 2 days at 37°C. Samples were retrieved, washed twice in PBS containing 0.2% Tween20 and 10μg/ml heparin (PTwH) and incubated with primary antibody solution (Iba1 1:500; PTwH/5%DMSO/3% goat serum) for 7 days at 37°C. After several washes, samples were incubated with secondary antibody solution (1:500 in PTwH/3% goat serum) for 4 days at 37°C. Prior to clearing, the samples were washed in PTwH and embedded in 2% Phytagel (Sigma Aldrich #P8169) in water. The embedded tissue was then dehydrated using an ascending series of Methanol/PBS and incubated overnight incubation in a mixture of 33% dichloromethan (DCM) and 66% MeOH at RT. Samples were further delipidated by incubation in 100% DCM for 40min and transferred to pure ethyl cinnamate (Eci; Sigma Aldrich #112372) as clearing reagent. Tissues became transparent after 15min in Eci and were stored at RT until imaging. Light sheet microscopy was performed using a LaVision Ultramicroscope II equipped with 2x objective, corrected dipping cap and zoom body. Spinal cords were mounted onto the sample holder with the dorsal/ventral axis facing down (z imaging axis = dorsoventral axis spinal cord). The holder was placed into the imaging chamber filled with Eci. Images were acquired in 3D multicolour mode with the following specifications: 5μm sheet thickness; 40% sheet width; 2x zoom; 4μm z-step size; one site sheet illumination; 100ms camera exposure time; full field of view. Autofluorescence was recorded using 488nm laser excitation (80% laser power) and a 525/40 emmision filter and red fluorescence was recorded using 561nm laser excitation (30% laser power) and 585/40 emission filters. Images were loaded into Vision4D 3.0 (Arivis) and the image set was cropped to 500 – 2000 pixels corresponding to 2.2 mm of spinal cord length. The volume of the spinal cord was determined by performing an automatic intensity thresholding on the autofluorescence channel. Phagocytes were detected by running a manual intensity thresholding on the 561nm channel and Iba1 cell accumulation with a volume of <1000μm3 was considered lesion positive. Then total lesion volume as well as the lesion volume fraction in respect to the total spinal cord volume were calculated. For 3D rendering, the autofluorescence and Iba1 channel were depicted in green and red pseudocolor, respectively. High-resolution images as well as videos were created using the Arivis 4D viewer.

### Laser capture microdissection

Serial cryostat coronal brain sections at the level of Bregma 1 mm to 0.5 mm were prepared (Leica) for single cell isolation). 700 and 800 EYFP+ neurons for each individual sample were microdissected (Arcturus Veritas microdissection system with fluorescence package) from the motor and somatosensory cortex and captured in HS Transfer Cap (Molecular Devices). Cells were only collected if no adjacent nuclei were detected in close proximity. Successful cutting and collection steps were subsequently validated in bright-field and fluorescent mode on the quality control slot of the device. Microdissected cells were lysed in 100 μl of RNA lysis buffer (Qiagen, Hilden, Germany) and stored at −80°C until further use. All procedures were done under RNase-free conditions.

### Microarray expression analysis

Total RNA of pooled single cells was resuspended with pretested T7-tagged dT21V oligonucleotides. Two-round T7-RNA polymerase-mediated linear amplification was performed according to optimized protocols for low-input RNA amounts (Small Sample Target Labeling Assay Version II, Affymetrix). Biotin-labeled second-round aRNA was generated with an NTP-mix containing Biotin-11-CTP and Biotin-16-UTP (PerkinElmer, Boston, MA) at 2 mM. Biotin-labeled amplified RNA (aRNA) size distribution and quantity was analyzed with the Agilent 2100 Bioanalyser using the RNA 6000 Nano LabChip kit (Agilent Technologies, Boeblingen, Germany). Samples with lower size compressed RNA products were discarded. At least 5 μg of labeled cRNA was fragmented by heating the sample to 95°C for 35 min in a volume of 20 μl containing 40 mM Tris acetate pH 8.1, 100 mM KOAc, and 30 mM MgOAc. Fragmentation was checked by alkaline agarose electrophoresis. Hybridization, washing, staining, and scanning were performed under standard conditions as described by the manufacturer. Mouse430A 2.0 genechips were used that contain over 22,600 probe sets representing transcripts and variants from over 14,000 mouse genes. Microarray raw data were exported using Gene chip operating software (Affymetrix). Normalization and higher-level analysis were done in R (for packages see Key Resources table). Normalization was carried out using the Robust Multichip Average (RMA) model implemented in the R package Affy at default settings. The normalized microarray data was quality controlled (box-plot analysis, principal component analysis, and Spearman correlation tree) which led to the exclusion of two microarrays. The remaining data were re-normalized, log transformed and filtered based on absolute expression values (100 fold changed signal intensity cutoff). Probe sets with a fold change higher than 1.5 were included in further analysis for single gene analysis. A fold change threshold of 1.3 was applied for further pathway level analysis using gene set enrichment analysis (GSEA; https://www.broadinstitute.org/gsea/). GSEA was performed with 5214 different gene sets obtained from Molecular Signature Database (MSigDB) at the Broad Institute (MIT).

### Human single-nuclei transcriptome sequencing datasets

Human single-nuclei RNA sequencing profiles were obtained from two available datasets, GSE118257 (Jäkel et al. 2019) and GSE124335 (Schirmer et al. 2019), and re-analyzed by R package Seurat v3.2.3. Both datasets were filtered and embedded according to parameters of original publications. Annotations of neurons, oligodendrocytes, and astrocytes were confirmed using marker gene expression of the different cell types (Figure S3). Subsequently, gene counts from neuron, oligodendrocyte and astrocyte subsets from both datasets were merged by applying Canonical Correlation Analysis (CCA) integration method. Uniform Manifold Approximation and Projection (UMAP) was used to visualize cell merging results. For each cell type, pairwise comparisons (MS lesion versus control and MS non-lesion versus control) of expression of genes related to cholesterol synthesis and metabolism were computed using normalized gene counts by Model-based Analysis of Single-cell Transcriptomics (MAST) R package v.1.12.0. Heatmap visualization was computed using R package pheatmap v.1.0.12 (Pretty Heatmaps).

### Spinal cord co-cultures

Spinal cord co-cultures were established as described (Bijland et al., 2019) from embryonic day 13 mouse embryos. Cells were plated initially in 12.5 % horse serum and fed the following day and every second or third day thereafter with serum-free differentiation medium. On day in vitro (DIV) 14 (before myelination commences around DIV 17), cultures were treated with cholesterol (10 μg/ml), cholesterol and tetrodotoxin (TTX) (1μM, Tocris), or 0.1% ethanol (vehicle control) for 3 days. On DIV 17, bromodeoxyuridine (BrdU; 10 μM) was added to the cultures for 2 hours. Cultures were fixed with PFA and washed in PBS. Cells were permeabilized for 10 minutes in 0.5% Triton X in PBS and incubated for 48 hours at 4°C in rabbit anti-NG2 (AB5320, Merck Millipore; 1:500) and rat anti-MBP (MCA409, Biorad, 1:500) in 10 % goat serum, 1 % bovine serum albumin in PBS. Following application of Alexa 596 anti-rabbit IgG and Alexa 488 anti-rat IgG (Invitrogen, 1:1000 for 1 hour), cells were fixed in 50:50 acetic acid and ethanol for 10 minutes and DNA was denatured in 2M HCl for 30 minutes. Then anti-BrdU (MCA2483T, Biorad; 1:500) was added in blocking buffer and incubated overnight. Alexa 647 anti-mouse IgG1 (Invitrogen, 1:1000) was added for ~1 hour at room temperature and coverslips were mounted in Mowiol with DAPI (2.5 μg/ml). 10 predefined locations were selected in the DAPI channel, and images were captured at 10x magnification using a Zeiss Axio Imager M.2 with an AxioCam MRm. Analysis of MBP positive area was performed by automated thresholding (Triangle, Image J). Quantification of the number of NG2 and BrdU double positive cells was done on binarized images and normalized to the DAPI-positive cell density with the particle analyzer plugin (Image J).

### Whole cell current clamp and microelectrode arrays

Whole cell patch clamp recording was performed on DIV 21 cultures using an Axopatch 200B amplifier with a Digidata 1440A digital acquisition system and pClamp 10 software (Molecular Devices). Experiments were performed at 37°C in atmospheric air using an extracellular solution containing (in mM): 144 NaCl, 5.3 KCl, 2.5 CaCl_2_, 1 MgCl_2_, 10 HEPES, 10 mM glucose, pH 7.4. The pipette solution contained (in mM): 130 mM K^+^ gluconate, 4 mM NaCl, 0.5 mM CaCl2, 10 mM HEPES, 0.5 EGTA pH 7.2. Borosilicate glass pipettes were pulled to a resistance of 3-8 MΩ. Liquid junction potentials were measured as 20 mV and traces were offset by this value. For microelectrode arrays, cultures were plated and maintained on commercial MEAs (60MEA200/30iR-Ti-gr; Multi Channel Systems, Reutlingen, Germany) as described previously (Bijland *et al.*, 2019). A fluorinated ethylene-propylene membrane (ALA MEA-MEM-SHEET) sealed the MEA culture dishes. The recordings were done in differentiation medium. Signals were digitally filtered at 3 Hz high pass filter, 1 kHz low pass filter and amplified up to x20,000. A digital notch filter was used to remove 60 Hz noise during recording. For data acquisition and analysis, spikes and potentials were sorted and counted from 3-minute gap-free recordings using the pCLAMP10 software (Molecular Devices Corporation, California, USA).

## QUANTIFICATION AND STATISTICAL ANALYSIS

Number of animals for each experiment is provided in the figure legends. No statistical methods were used to pre-determine sample sizes but our sample sizes are similar to those reported in previous publications (Berghoff *et al.*, 2021). No inclusion or exclusion criteria were used if not otherwise stated. Studies were conducted blinded to investigators and/or formally randomized. Data are expressed as mean ± SEM unless otherwise indicated. For statistical analysis, unpaired two-sided Student’s t-test, one-way ANOVA or two-way ANOVA with Sidak’s or Tukey’s post tests were applied. Normality was tested by using the Kolmogorov-Smirnov test. If the n was below 5, we assumed normal distribution. “Signal-to-Noise ratio” (SNR) statistics were used to rank genes for GSEA of microarray data. Linear model fitting and subsequent testing for differential expression by empirical Bayes variance moderation method implemented in R packaged limma v3.42.2 was applied to the 6-month neuron microarray data. Wilcoxon Rans Sum test was used for analysis of snRNAseq data material. Data analysis was performed using GraphPad Prism Software Version 6 (GraphPad) and the R software. A value of p<0.05 was considered statistically significant. Asterisks depict statistically significant differences (* p<0.05, ** p<0.01, *** p<0.001).

## Acknowledgments

We cordially thank Annette Fahrenholz and Tanja Freerck for technical support. We thank Charles Stiles, John Alberta, Said Ghandour for generous gifts of antibodies. This work was funded by the Deutsche Forschungsgemeinschaft (SA 2014/2-1 to GS), the UK MS Society (Grant 127 to JE); Medical Research Scotland (PhD studentship 791-2014 to EB).

## Author Contributions

SAB and GS planned and designed the study. SAB and LS were involved in all experiments. TS, YZ, and SBo performed reanalysis of human snRNAseq datasets. LH and FO did flow cytometry. TI and PS performed lipid mass spectrometry. MHV and AMS performed histology. CD and AOS did light sheet microscopy. DKB was involved in behavior experiments. SW, KAN, and MR analyzed genetic myelin mutants. JME conducted myelinating cell culture experiments. KM and EB conducted electrophysiology experiments on myelinating cell cultures. SAB and GS wrote and edited the manuscript. All authors approved the manuscript.

## LEAD CONTACT AND MATERIAL AVAILABILITY

Further information and requests for resources and reagents should be directed to and will be fulfilled by the Lead Contact, Gesine Saher. This study did not generate new unique reagents.

## References

Almeida, R.G., Williamson, J.M., Madden, M.E., Early, J.J., Voas, M.G., Talbot, W.S., Bianco, I.H., and Lyons, D.A. (2020). Synaptic vesicle fusion along axons is driven by myelination and subsequently accelerates sheath growth in an activity-regulated manner. bioRxiv, 2020.2008.2028.271593. 10.1101/2020.08.28.271593.

Bacmeister, C.M., Barr, H.J., McClain, C.R., Thornton, M.A., Nettles, D., Welle, C.G., and Hughes, E.G. (2020). Motor learning promotes remyelination via new and surviving oligodendrocytes. Nature neuroscience 23, 819–831. 10.1038/s41593-020-0637-3.

Berghoff, S.A., Duking, T., Spieth, L., Winchenbach, J., Stumpf, S.K., Gerndt, N., Kusch, K., Ruhwedel, T., Mobius, W., and Saher, G. (2017a). Blood-brain barrier hyperpermeability precedes demyelination in the cuprizone model. Acta Neuropathol Commun 5, 94. 10.1186/s40478-017-0497-6.

Berghoff, S.A., Gerndt, N., Winchenbach, J., Stumpf, S.K., Hosang, L., Odoardi, F., Ruhwedel, T., Böhler, C., Barrette, B., Stassart, R., et al. (2017b). Dietary cholesterol promotes repair of demyelinated lesions in the adult brain. Nature Communications 8, 14241. 10.1038/ncomms14241.

Berghoff, S.A., Spieth, L., Sun, T., Hosang, L., Schlaphoff, L., Depp, C., Duking, T., Winchenbach, J., Neuber, J., Ewers, D., et al. (2021). Microglia facilitate repair of demyelinated lesions via post-squalene sterol synthesis. Nat Neurosci 24, 47–60. 10.1038/s41593-020-00757-6.

Bijland, S., Thomson, G., Euston, M., Michail, K., Thummler, K., Mucklisch, S., Crawford, C.L., Barnett, S.C., McLaughlin, M., Anderson, T.J., et al. (2019). An in vitro model for studying CNS white matter: functional properties and experimental approaches. F1000Res 8, 117. 10.12688/f1000research.16802.1.

Borggrewe, M., Grit, C., Vainchtein, I.D., Brouwer, N., Wesseling, E.M., Laman, J.D., Eggen, B.J.L., Kooistra, S.M., and Boddeke, E. (2021). Regionally diverse astrocyte subtypes and their heterogeneous response to EAE. Glia 69, 1140–1154. 10.1002/glia.23954.

Camargo, N., Brouwers, J.F., Loos, M., Gutmann, D.H., Smit, A.B., and Verheijen, M.H. (2012). High-fat diet ameliorates neurological deficits caused by defective astrocyte lipid metabolism. FASEB journal : official publication of the Federation of American Societies for Experimental Biology 26, 4302–4315. 10.1096/fj.12-205807.

Cantoni, C., Bollman, B., Licastro, D., Xie, M., Mikesell, R., Schmidt, R., Yuede, C.M., Galimberti, D., Olivecrona, G., Klein, R.S., et al. (2015). TREM2 regulates microglial cell activation in response to demyelination in vivo. Acta Neuropathol 129, 429–447. 10.1007/s00401-015-1388-1.

Crawford, D.K., Mangiardi, M., Xia, X., Lopez-Valdes, H.E., and Tiwari-Woodruff, S.K. (2009). Functional recovery of callosal axons following demyelination: a critical window. Neuroscience 164, 1407–1421. 10.1016/j.neuroscience.2009.09.069.

Cunha, M.I., Su, M., Cantuti-Castelvetri, L., Muller, S.A., Schifferer, M., Djannatian, M., Alexopoulos, I., van der Meer, F., Winkler, A., van Ham, T.J., et al. (2020). Pro-inflammatory activation following demyelination is required for myelin clearance and oligodendrogenesis. J Exp Med 217. 10.1084/jem.20191390.

Demerens, C., Stankoff, B., Logak, M., Anglade, P., Allinquant, B., Couraud, F., Zalc, B., and Lubetzki, C. (1996). Induction of myelination in the central nervous system by electrical activity. Proceedings of the National Academy of Sciences of the United States of America 93, 9887–9892. 10.1073/pnas.93.18.9887.

Dietschy, J.M. (2009). Central nervous system: cholesterol turnover, brain development and neurodegeneration. Biological chemistry 390, 287–293. 10.1515/BC.2009.035.

Dietschy, J.M., and Turley, S.D. (2004). Thematic review series: brain Lipids. Cholesterol metabolism in the central nervous system during early development and in the mature animal. Journal of lipid research 45, 1375–1397. 10.1194/jlr.R400004-JLR200.

Edgar, J.M., McLaughlin, M., Werner, H.B., McCulloch, M.C., Barrie, J.A., Brown, A., Faichney, A.B., Snaidero, N., Nave, K.A., and Griffiths, I.R. (2009). Early ultrastructural defects of axons and axon-glia junctions in mice lacking expression of Cnp1. Glia 57, 1815–1824. 10.1002/glia.20893.

Edgar, J.M., McLaughlin, M., Yool, D., Zhang, S.C., Fowler, J.H., Montague, P., Barrie, J.A., McCulloch, M.C., Duncan, I.D., Garbern, J., et al. (2004). Oligodendroglial modulation of fast axonal transport in a mouse model of hereditary spastic paraplegia. J Cell Biol 166, 121–131. 10.1083/jcb.200312012.

Franklin, R.J.M., Frisen, J., and Lyons, D.A. (2020). Revisiting remyelination: Towards a consensus on the regeneration of CNS myelin. Semin Cell Dev Biol. 10.1016/j.semcdb.2020.09.009.

Fünfschilling, U., Jockusch, W.J., Sivakumar, N., Möbius, W., Corthals, K., Li, S., Quintes, S., Kim, Y., Schaap, I.A., Rhee, J.S., et al. (2012). Critical time window of neuronal cholesterol synthesis during neurite outgrowth. The Journal of neuroscience : the official journal of the Society for Neuroscience 32, 7632–7645. 10.1523/JNEUROSCI.1352-11.2012.

Gibson, E.M., Purger, D., Mount, C.W., Goldstein, A.K., Lin, G.L., Wood, L.S., Inema, I., Miller, S.E., Bieri, G., Zuchero, J.B., et al. (2014). Neuronal activity promotes oligodendrogenesis and adaptive myelination in the mammalian brain. Science 344, 1252304. 10.1126/science.1252304.

Hess, K., Starost, L., Kieran, N.W., Thomas, C., Vincenten, M.C.J., Antel, J., Martino, G., Huitinga, I., Healy, L., and Kuhlmann, T. (2020). Lesion stage-dependent causes for impaired remyelination in MS. Acta Neuropathol 140, 359–375. 10.1007/s00401-020-02189-9.

Ioannou, M.S., Jackson, J., Sheu, S.H., Chang, C.L., Weigel, A.V., Liu, H., Pasolli, H.A., Xu, C.S., Pang, S., Matthies, D., et al. (2019). Neuron-Astrocyte Metabolic Coupling Protects against Activity-Induced Fatty Acid Toxicity. Cell 177, 1522–1535 e1514. 10.1016/j.cell.2019.04.001.

Itoh, N., Itoh, Y., Tassoni, A., Ren, E., Kaito, M., Ohno, A., Ao, Y., Farkhondeh, V., Johnsonbaugh, H., Burda, J., et al. (2018). Cell-specific and region-specific transcriptomics in the multiple sclerosis model: Focus on astrocytes. Proc Natl Acad Sci U S A 115, E302–E309. 10.1073/pnas.1716032115.

Jurevics, H., Largent, C., Hostettler, J., Sammond, D.W., Matsushima, G.K., Kleindienst, A., Toews, A.D., and Morell, P. (2002). Alterations in metabolism and gene expression in brain regions during cuprizone-induced demyelination and remyelination. J Neurochem 82, 126–136.

Kassmann, C.M., Lappe-Siefke, C., Baes, M., Brugger, B., Mildner, A., Werner, H.B., Natt, O., Michaelis, T., Prinz, M., Frahm, J., and Nave, K.A. (2007). Axonal loss and neuroinflammation caused by peroxisome-deficient oligodendrocytes. Nature genetics 39, 969–976. 10.1038/ng2070.

Korinek, M., Gonzalez-Gonzalez, I.M., Smejkalova, T., Hajdukovic, D., Skrenkova, K., Krusek, J., Horak, M., and Vyklicky, L. (2020). Cholesterol modulates presynaptic and postsynaptic properties of excitatory synaptic transmission. Sci Rep 10, 12651. 10.1038/s41598-020-69454-5.

Lappe-Siefke, C., Goebbels, S., Gravel, M., Nicksch, E., Lee, J., Braun, P.E., Griffiths, I.R., and Nave, K.A. (2003). Disruption of Cnp1 uncouples oligodendroglial functions in axonal support and myelination. Nat Genet 33, 366–374. 10.1038/ng1095.

Linetti, A., Fratangeli, A., Taverna, E., Valnegri, P., Francolini, M., Cappello, V., Matteoli, M., Passafaro, M., and Rosa, P. (2010). Cholesterol reduction impairs exocytosis of synaptic vesicles. J Cell Sci 123, 595–605. 10.1242/jcs.060681.

Marisca, R., Hoche, T., Agirre, E., Hoodless, L.J., Barkey, W., Auer, F., Castelo-Branco, G., and Czopka, T. (2020). Functionally distinct subgroups of oligodendrocyte precursor cells integrate neural activity and execute myelin formation. Nature neuroscience 23, 363–374. 10.1038/s41593-019-0581-2.

Mathews, E.S., and Appel, B. (2016). Cholesterol Biosynthesis Supports Myelin Gene Expression and Axon Ensheathment through Modulation of P13K/Akt/mTor Signaling. Journal of Neuroscience 36, 7628–7639. 10.1523/Jneurosci.0726-16.2016.

Mauch, D.H., Nagler, K., Schumacher, S., Goritz, C., Muller, E.C., Otto, A., and Pfrieger, F.W. (2001). CNS synaptogenesis promoted by glia-derived cholesterol. Science 294, 1354–1357. 10.1126/science.294.5545.1354.

Miron, V.E., Boyd, A., Zhao, J.W., Yuen, T.J., Ruckh, J.M., Shadrach, J.L., van Wijngaarden, P., Wagers, A.J., Williams, A., Franklin, R.J.M., and Ffrench-Constant, C. (2013). M2 microglia and macrophages drive oligodendrocyte differentiation during CNS remyelination. Nature neuroscience 16, 1211–1218. 10.1038/nn.3469.

Miron, V.E., Rajasekharan, S., Jarjour, A.A., Zamvil, S.S., Kennedy, T.E., and Antel, J.P. (2007). Simvastatin regulates oligodendroglial process dynamics and survival. Glia 55, 130–143. 10.1002/glia.20441.

Miron, V.E., Zehntner, S.P., Kuhlmann, T., Ludwin, S.K., Owens, T., Kennedy, T.E., Bedell, B.J., and Antel, J.P. (2009). Statin therapy inhibits remyelination in the central nervous system. Am J Pathol 174, 1880–1890. 10.2353/ajpath.2009.080947.

Mitew, S., Gobius, I., Fenlon, L.R., McDougall, S.J., Hawkes, D., Xing, Y.L., Bujalka, H., Gundlach, A.L., Richards, L.J., Kilpatrick, T.J., et al. (2018). Pharmacogenetic stimulation of neuronal activity increases myelination in an axon-specific manner. Nature communications 9, 306. 10.1038/s41467-017-02719-2.

Nikic, I., Merkler, D., Sorbara, C., Brinkoetter, M., Kreutzfeldt, M., Bareyre, F.M., Bruck, W., Bishop, D., Misgeld, T., and Kerschensteiner, M. (2011). A reversible form of axon damage in experimental autoimmune encephalomyelitis and multiple sclerosis. Nature medicine 17, 495–499. 10.1038/nm.2324.

Ortiz, F.C., Habermacher, C., Graciarena, M., Houry, P.Y., Nishiyama, A., Nait Oumesmar, B., and Angulo, M.C. (2019). Neuronal activity in vivo enhances functional myelin repair. JCI Insight 5. 10.1172/jci.insight.123434.

Pfeiffer, F., Frommer-Kaestle, G., and Fallier-Becker, P. (2019). Structural adaption of axons during de- and remyelination in the Cuprizone mouse model. Brain pathology 29, 675–692. 10.1111/bpa.12748.

Plemel, J.R., Liu, W.Q., and Yong, V.W. (2017). Remyelination therapies: a new direction and challenge in multiple sclerosis. Nature reviews. Drug discovery 16, 617–634. 10.1038/nrd.2017.115.

Reich, D.S., Lucchinetti, C.F., and Calabresi, P.A. (2018). Multiple Sclerosis. The New England journal of medicine 378, 169–180. 10.1056/NEJMra1401483.

Saab, A.S., and Nave, K.A. (2017). Myelin dynamics: protecting and shaping neuronal functions. Current opinion in neurobiology 47, 104–112. 10.1016/j.conb.2017.09.013.

Saher, G., Brugger, B., Lappe-Siefke, C., Mobius, W., Tozawa, R., Wehr, M.C., Wieland, F., Ishibashi, S., and Nave, K.A. (2005). High cholesterol level is essential for myelin membrane growth. Nat Neurosci 8, 468–475. 10.1038/nn1426.

Saher, G., Rudolphi, F., Corthals, K., Ruhwedel, T., Schmidt, K.F., Lowel, S., Dibaj, P., Barrette, B., Mobius, W., and Nave, K.A. (2012). Therapy of Pelizaeus-Merzbacher disease in mice by feeding a cholesterol-enriched diet. Nat Med 18, 1130–1135. 10.1038/nm.2833.

Scalfari, A. (2019). MS progression is predominantly driven by age-related mechanisms – YES. Multiple sclerosis 25, 902–904. 10.1177/1352458518820633.

Stassart, R.M., Mobius, W., Nave, K.A., and Edgar, J.M. (2018). The Axon-Myelin Unit in Development and Degenerative Disease. Frontiers in neuroscience 12, 467. 10.3389/fnins.2018.00467.

Thelen, K.M., Falkai, P., Bayer, T.A., and Lutjohann, D. (2006). Cholesterol synthesis rate in human hippocampus declines with aging. Neurosci Lett 403, 15–19. 10.1016/j.neulet.2006.04.034.

Thiele, C., Hannah, M.J., Fahrenholz, F., and Huttner, W.B. (2000). Cholesterol binds to synaptophysin and is required for biogenesis of synaptic vesicles. Nat Cell Biol 2, 42–49. 10.1038/71366.

Trevisiol, A., Kusch, K., Steyer, A.M., Gregor, I., Nardis, C., Winkler, U., Kohler, S., Restrepo, A., Mobius, W., Werner, H.B., et al. (2020). Structural myelin defects are associated with low axonal ATP levels but rapid recovery from energy deprivation in a mouse model of spastic paraplegia. PLoS biology 18, e3000943. 10.1371/journal.pbio.3000943.

Vance, J.E., Campenot, R.B., and Vance, D.E. (2000). The synthesis and transport of lipids for axonal growth and nerve regeneration. Biochimica et biophysica acta 1486, 84–96. 10.1016/s1388-1981(00)00050-0.

Voskuhl, R.R., Itoh, N., Tassoni, A., Matsukawa, M.A., Ren, E., Tse, V., Jang, E., Suen, T.T., and Itoh, Y. (2019). Gene expression in oligodendrocytes during remyelination reveals cholesterol homeostasis as a therapeutic target in multiple sclerosis. Proc Natl Acad Sci U S A 116, 10130–10139. 10.1073/pnas.1821306116.

Wehr, M.C., Laage, R., Bolz, U., Fischer, T.M., Grunewald, S., Scheek, S., Bach, A., Nave, K.A., and Rossner, M.J. (2006). Monitoring regulated protein-protein interactions using split TEV. Nature methods 3, 985–993. 10.1038/nmeth967.

Xu, Q., Bernardo, A., Walker, D., Kanegawa, T., Mahley, R.W., and Huang, Y. (2006). Profile and regulation of apolipoprotein E (ApoE) expression in the CNS in mice with targeting of green fluorescent protein gene to the ApoE locus. The Journal of neuroscience : the official journal of the Society for Neuroscience 26, 4985–4994. 10.1523/JNEUROSCI.5476-05.2006.

Zhao, C., Deng, Y., Liu, L., Yu, K., Zhang, L., Wang, H., He, X., Wang, J., Lu, C., Wu, L.N., et al. (2016). Dual regulatory switch through interactions of Tcf7l2/Tcf4 with stage-specific partners propels oligodendroglial maturation. Nat Commun 7, 10883. 10.1038/ncomms10883.

