## Supplementary figures for "Neuronal cholesterol synthesis is essential for repair of chronically demyelinated lesions in mice"

**Supplement**

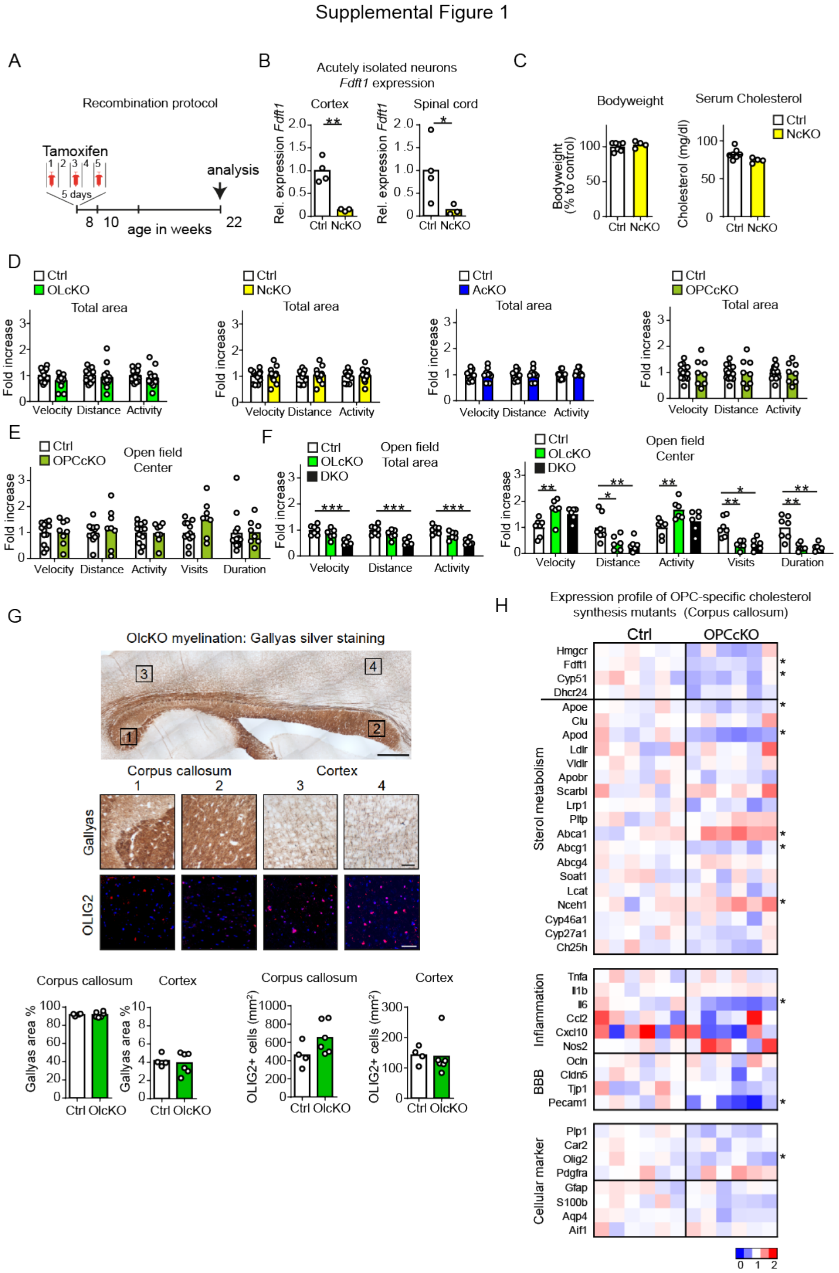


**Figure S1. Genetic inactivation of *Fdft1* in neurons, oligodendrocytes and astrocytes, Related to figure 1**

(A) Scheme depicting tamoxifen induced recombination protocol used for conditional inactivation of *Fdft1* in oligodendrocytes and astrocytes.

(B) Expression of *Fdft1* determined by RT-qPCR in isolated neurons from cortex and spinal cord of NcKO animals normalized to controls (two-sided Student's t-test).

(C) Mean body weight and total serum cholesterol with individual values of neuronal cholesterol mutants (NcKO, n=4) and controls (n=7) at the age of 22 weeks (two-sided Student's t-test).

(D) Open field test (total area data) of conditional cholesterol mutants in oligodendrocytes (OLcKO; n=14), neurons (NcKO; n=11), astrocytes (AcKO; n=9) and oligodendrocyte precursor cells (OPCcKO) compared to corresponding controls (n=8-12). Data are shown as mean with values of individual animals.

(E) Open field test (center data) of OPCcKO mutants compared to corresponding controls (n=8-12).

(F) Open field test (total area and center data) of OLcKO and OPC/oligodendrocyte double knockouts (DKO) of cholesterol synthesis compared to corresponding controls (n=6-7; one-way ANOVA with Sidak's post-test).

(G) Representative Gallyas staining to visualize myelin in brain sections of OLcKO mice at the age of 22 weeks (scale 500 µm) with details of boxed areas below showing Gallyas and Olig2 immunolabeling (scale 50 µm). Quantification of myelination (Gallyas) and Olig2 (oligodendrocyte linage cells) was done in the corpus callosum and cortex.

(H) Gene expression profile of genes related to cholesterol metabolism, inflammation, blood-brain barrier and cellular identity in corpus callosum samples of OPCcKO mutants and controls. Heat maps show fold expression normalized to controls. Each square represents a biological replicate (n=6, two-sided Student's t-test).

***p<0.001, **p<0.01, *p<0.05

Supplemental Figure 2

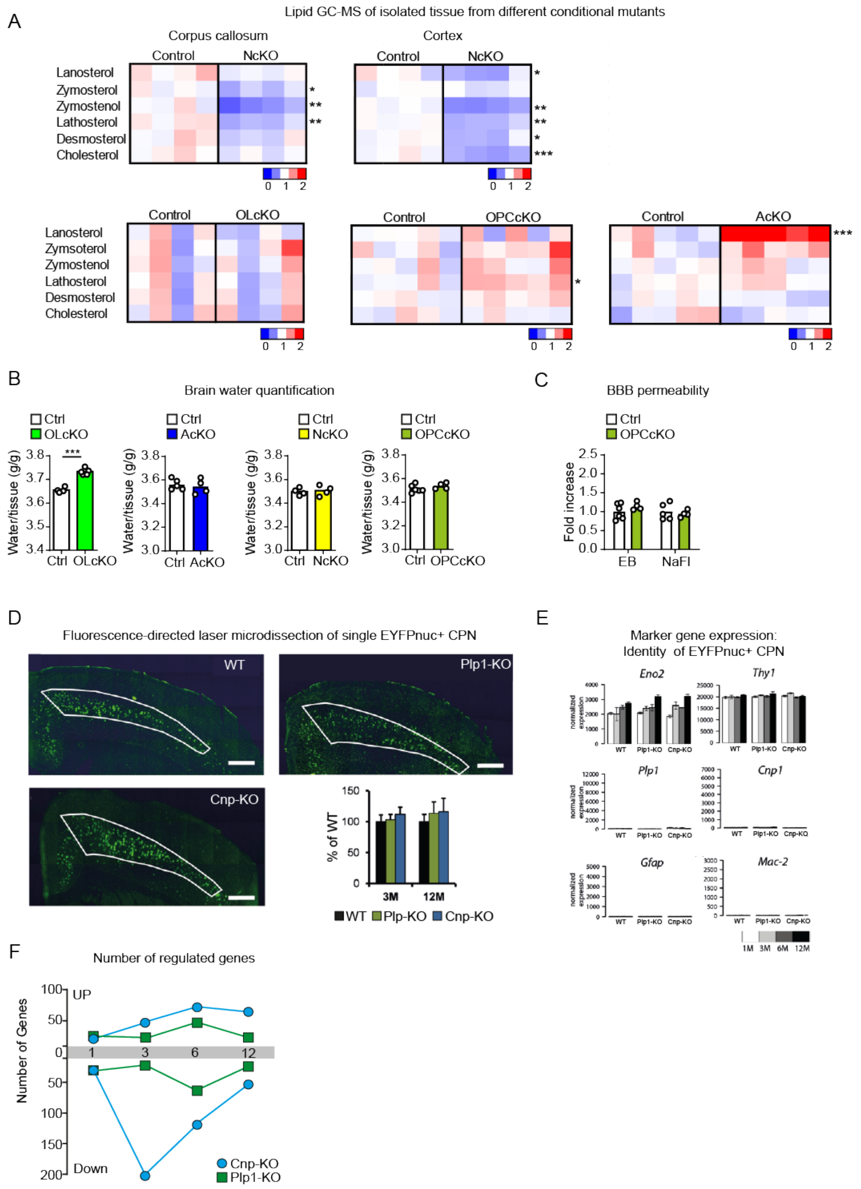


**Figure S2. Isolation of EYFPnuc+ neurons from CNP and PLP mutants, Related to figure 1 and 2**
(A) Relative abundance of sterol intermediates in isolated tissue from conditional mutants (n=4-5) compared to untreated controls (n=4; set to 1). Corpus callosum and cortical tissue was isolated from NcKO (n=4) animals and corresponding controls (n=4). Thalamic samples of OlcKO (n=4), OPCcKO (n=5) and AcKO (n=5) and respective controls (n=4-5) were measured by GC-MS (two-sided Student's t-test). One square represents an animal.
(B) Brain water content in OLcKO (n=5), AcKO (n=5), NcKO (n=4) and OPCcKO (n=4) mutants compared to corresponding controls (n=5, two-sided Student's t-test, ***p<0.001, **p<0.01).
(C) Extravasation of EB and NaFl in OPCcKO (n=4) compared to controls (n=5).
(D) Representative sections of cerebral cortex of wild type and CNP and PLP mutant mice that additionally harbor the EYFPnuc transgene (scale 50 µm). Mean density of EYFPnuc+ CPN cells ± SD in motor and somatosensory cortex (outlined area) in CNP and PLP mutant mice at the age of 3 and 12 months relative to wild type mice (n=3).
(E) Validation of the cell type identity of laser microdissected cells by marker gene expression using the neuronal markers *Eno2* and *Thy1*, oligodendrocyte-specific marker genes *Plp1* and *Cnp1*, the astrocyte marker *Gfap*, and the microglial marker Mac-2 (*Lgals*). Bar plots represent arithmetic means (n=3) ± SD.
(F) Number of differentially expressed genes in CPN of CNP and PLP mutants.

Supplemental Figure 3

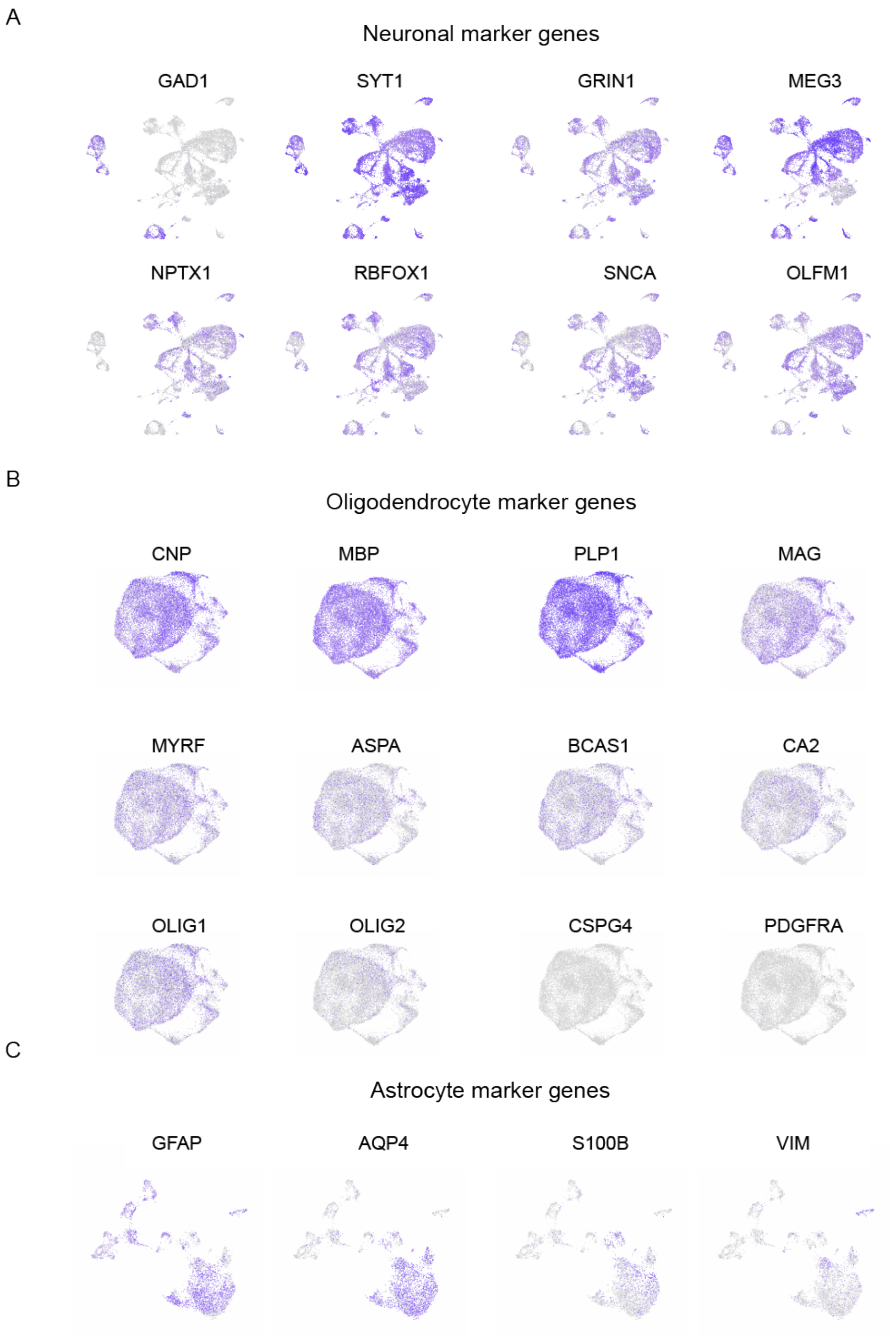


**Figure S3. Re-analysis of human snRNAseq MS datasets, Related to figure 3**
(A) Expression of selected marker genes in the neuronal subset (*GAD1, SYT1, GRIN1, MEG3, NPTX1, RBFOX1, SNCA, OLFM1*) of snRNAseq datasets.
(B) Expression of selected oligodendrocyte marker genes (*CNP, MBP, PLP1, MAG, MYRF, ASPA, BCAS1, CA2*), oligodendrocyte linage marker genes (*OLIG1, OLIG2*) and OPC marker genes (*CSPG4, PDGFRA*) in oligodendrocytes of snRNAseq datasets.
(C) Expression of selected marker genes in astrocytes (*GFAP, AQP4, S100B, VIM*) of snRNAseq datasets.

Supplemental Figure 4

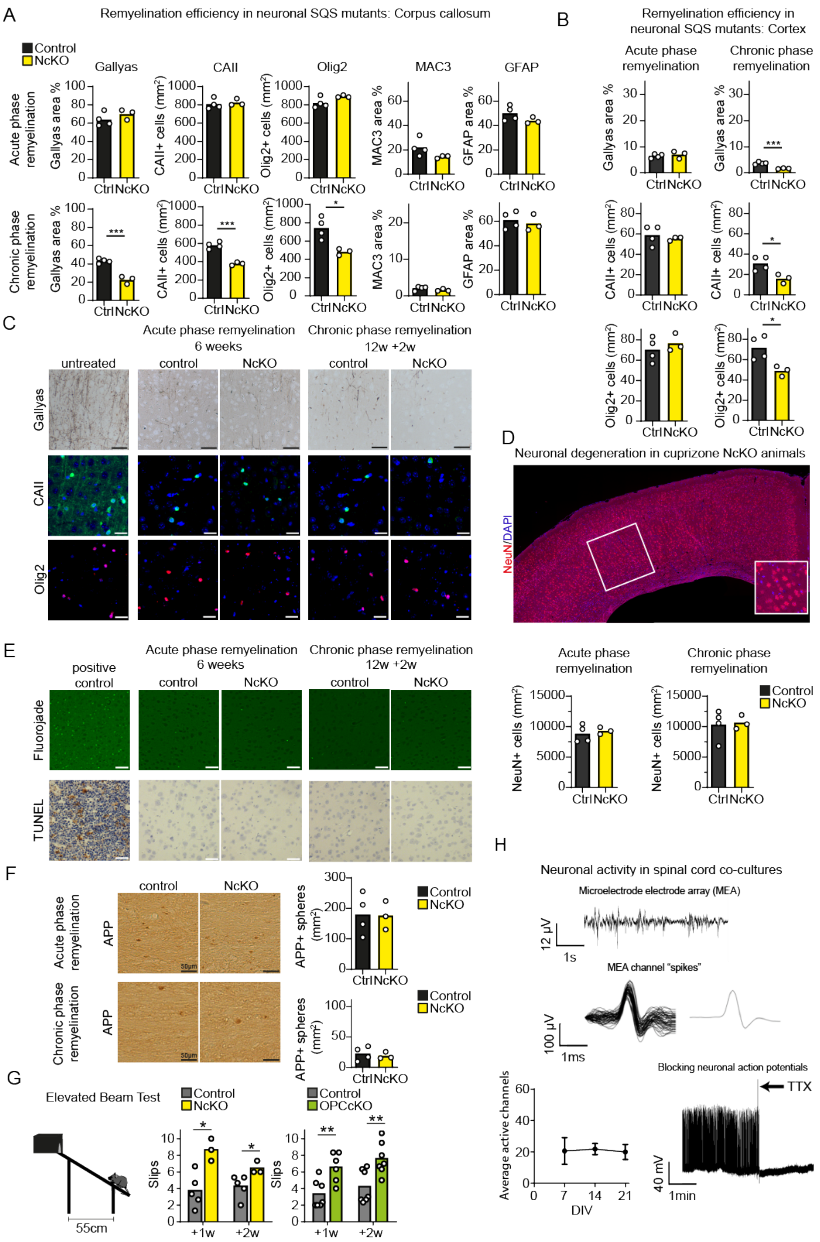


**Figure S4. Remyelination efficiency following neuronal cholesterol synthesis ablation, Related to figure 4**

(A-B) Quantification of histochemical stainings in the corpus callosum (A) and cortex (B) of NcKO mutants (n=3) and controls (n=4) in acute-phase remyelination and chronic-phase remyelination, evaluating myelination (Gallyas), density of oligodendrocytes (CAII), density of oligodendrocyte lineage cells (Olig2), microgliosis (MAC3) and astrogliosis (GFAP). Data represent the mean with individual values (two-sided Student's t-test).

(C) Representative cortical images of histological stainings of untreated, control and NcKO mice, detecting myelin (Gallyas), oligodendrocytes (CAII), and oligodendrocyte lineage cells (Olig2) (scale 50 µm).

(D) Representative micrograph of NeuN+ neuronal cell bodies (scale 500 µm) with quantification of neuron density in NcKO and control mice during acute-phase and chronic-phase remyelination below. The box area (cortical layer 4-6) indicates the area used for quantification.

(E) Apoptotic cell labeling of cortical samples from NcKO and control animals following acute and chronic cuprizone using Fluorojade and TUNEL staining (scale 50 µm). Diphtheria toxin mediated neuronal degeneration within cortical tissue (Fluorojade) and embryonic liver sections (TUNEL) were used as positive controls.

(F) Representative micrographs of APP labeling in the corpus callosum from NcKO and control animals during acute-phase and chronic-phase remyelination with quantification (scale 50 µm).

***$p<0.001$, *$p<0.05$

(G) Elevated beam testing of NcKO (n=3) and OPCcKO (n=7) mutants compared to respective controls (n=5-6) during chronic remyelination (12+1w and 12+2w; one-way ANOVA with Sidak's post-test).

(H) (Upper) Representative extracellular recording from a microelectrode array at DIV 15 shows high frequency neuronal firing, presumably originating from several active neuronal elements. (Middle) Discrete spikes with a duration >1.5 ms and averaged trace from the microelectrode array (MEA), likely representing axonally-generated action potentials. (Lower left) Mean neuronal activity ± SEM quantified as proportion of active channels of the 59-electrode MEA, showing spontaneous electrical activity on ~30% of the channels for the whole culture period. (Lower right) Representative whole-cell recording in current-clamp mode at the neuronal resting potential (around -65 mV corrected for liquid junction potentials). The neuron shows a typical high frequency rate of spontaneous action potential firing (DIV 21). After addition of 1 µM TTX (arrow) neuronal action potentials are rapidly abolished.
